# Exact and approximate limit behaviour of the Yule tree’s cophenetic index

**DOI:** 10.1101/120931

**Authors:** Krzysztof Bartoszek

## Abstract

In this work we study the limit distribution of an appropriately normalized cophenetic index of the pure–birth tree conditioned on *n* contemporary tips. We show that this normalized phylogenetic balance index is a submartingale that converges almost surely and in *L*^2^. We link our work with studies on trees without branch lengths and show that in this case the limit distribution is a contraction–type distribution, similar to the Quicksort limit distribution. In the continuous branch case we suggest approximations to the limit distribution. We propose heuristic methods of simulating from these distributions and it may be observed that these algorithms result in reasonable tails. Therefore, we propose a way based on the quantiles of the derived distributions for hypothesis testing, whether an observed phylogenetic tree is consistent with the pure–birth process. Simulating a sample by the proposed heuristics is rapid, while exact simulation (simulating the tree and then calculating the index) is a time–consuming procedure. We conduct a power study to investigate how well the cophenetic indices detect deviations from the Yule tree and apply the methodology to empirical phylogenies.

## 1 Introduction

Phylogenetic trees are now a standard when analyzing groups of species. They are inferred from molecular sequences by algorithms that often assume a Markov chain for mutations at the individual positions of the genetic sequence (e.g. Ewens and Grant, 2005; Felsenstein, 2004; Yang, 2006). Given a phylogenetic tree it is often of interest to quantify the rate(s) of speciation and extinction for the studied species. To do this one commonly assumes a birth–death process with constant rates. However, the development of formal statistical tests whether a given tree comes from a given branching process model is an open area of research (see the still relevant “Work remaining” part at the end of Ch. 33 in Felsenstein, 2004). The reason for the apparent lack of widespread use of such tests (but see Blum and François, 2005) could be the lack of a commonly agreed on test statistic. This is as a tree is a complex object and there are multiple ways in which to summarize it in a single number.

One proposed way of summarizing a tree is through indices that quantify how balanced it is, i.e. how close is it to a fully symmetric tree. Two such indices have been with us for many years now: Sackin’s (Sackin, 1972) and Colless’ (Colless, 1982). Alternatively, McKenzie and Steel (2000) proposed to measure balance by counting cherries on the tree and they showed that after appropriate centring and scaling, this index converges to the standard normal distribution (for examples of other indices see Ch. 33 in Felsenstein, 2004).

Recently, a new balance index was proposed—the cophenetic index (Mir et al., 2013). The work here is inspired by private communication with evolutionary biologist Gabriel Yedid (current affiliation Nanjing Agricultural University, Nanjing, China) concerning the usage of the cophenetic index for significance testing of whether a given tree is consistent with the pure–birth process. He noticed that simulated distributions of the index have much heavier tails than those of the normal and t distributions and hence, comparing centred and scaled cophenetic indices with the usual Gaussian or t quantiles is not appropriate for significance testing. It would lead to a higher false positive rate—rejecting the null hypothesis of no extinction when a tree was generated by a pure–birth process.

Our aim here is to propose an approach for working analytically with the cophenetic index, especially to improve hypothesis tests for phylogenetic trees, i.e. how to recognize if the tree is out of the “Yule zone” (Yang et al., 2017). We show that there is a relationship between the cophenetic index and the Quicksort algorithm. This suggests that the methods exploring (e.g. Fill and Janson, 2000, 2001; Janson, 2015) the limiting distribution of the Quicksort algorithm can be an inspiration for studying analytical properties of the cophenetic index.

The paper is organized as follows. In Section 2 we formally define the cophenetic index (for trees with and without branch lengths) and present the most important results of the manuscript. We define an associated submartingale that converges almost surely and in *L*^2^ (Thm. 2.4), propose an elegant representation (Thm. 2.7) and a very promising approximation (Def. 2.8). Afterwards in Section 3, we show that in the discrete setting the limit law of the normalized cophenetic index is a contraction–type distribution. Based on this we propose alternative approximations to the limit law of the normalized (with branch lengths) cophenetic index. In Section 4 we describe heuristic algorithms to simulate from these limit laws, show simulated quantiles, explore the power of the cophenetic index to recognize deviations from the Yule tree (comparing with Sackin’s and Colless’ indices’ powers), and apply the indices to example empirical data. In Section 5 we prove the claims presented in Section 2 alongside other supporting results. Then, in Section 6 we study the second order properties of this decomposition and conjecture a Central Limit Theorem (CLT, Rem. 6.10). We end the paper with Section 7 by describing alternative representations of the cophenetic index.

## 2 The cophenetic index and summary of main results

Mir et al. (2013) recently proposed a new balance index for phylogenetic trees.

### Definition 2.1

**(Mir et al. (2013))** For a given phylogenetic tree on n tips and for each pair of tips (i,j) let 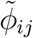 be the number of branches from the root to the most recent common ancestor of tips i and j. We then the define the discrete cophenetic index as

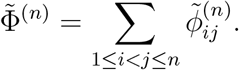

Mir et al. (2013) show that this index has a better resolution than the “traditional” ones. In particular the cophenetic index has a range of values of the order of O(n^3^) while Colless’ and Sackin’s ranges have an order of O(n^2^). Furthermore, unlike the other two previously mentioned, 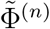 makes mathematical sense for trees that are not fully resolved (i.e. not binary).

In this work we study phylogenetic trees with branch lengths and hence consider a variation of the cophenetic index.

### Definition 2.2

For a given phylogenetic tree on n tips and for each pair of tips (i,j) let φ_ij_ be the time from the most recent common ancestor of tips i and j to the root/origin (depending on the tree model) of the tree. We then define the continuous cophenetic index as

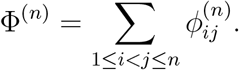

**Remark 2.3** In the original setting, when the distance between two nodes was measured by counting branches, Mir et al. (2013) did not consider the edge leading to the root. In our work here, where our prime concern is with trees with random branch lengths, we include the branch leading to the root. This is not a big difference, one just has to remember to add to each distance between nodes the same exponential (exp(1)—parametrization by the rate) random variable (see Section 5 for description of the tree’s growth).

The results of the present manuscript are built around a scaled version of the cophenetic index which is an almost surely and *L*^2^ convergent submartingale. We first introduce some notation. Let 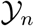 be the *σ*–algebra containing all the information on the Yule with *n* tips tree and define

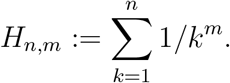

Below we present the main results concerning the cophenetic index, leaving the proofs and supporting theorems for Section 5.

### Theorem 2.4

Consider a scaled cophenetic index

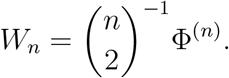

W_n_ is a positive submartingale that converges almost surely and in L^2^ to a finite first and second moment random variable.

### Definition 2.5

For k = 1,…, n−1 let us define 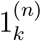 as the indicator random variable taking the value of 1 if a randomly sampled pair of species coalesced at the k–th (counting from the origin of the tree) speciation event.

We know (e.g. Bartoszek and Sagitov, 2015b; Stadler, 2009; Steel and McKenzie, 2001) that

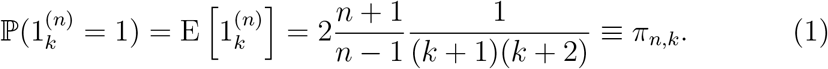

### Definition 2.6

For i = 1,…, n − 1 let us introduce the random variable

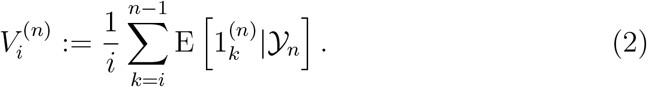

### Theorem 2.7

W_n_ can be represented as

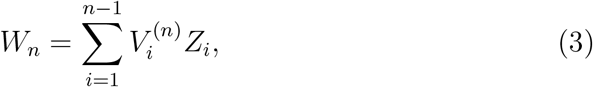

where Z_1_,…, Z_n−1_ are i.i.d. exp(1) random variables.

### Definition 2.8

Define the random variable 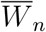 as

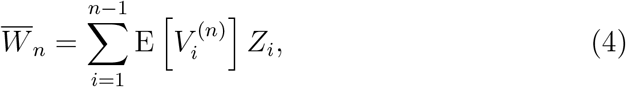

where Z_1_,…, Z_n−1_ are i.i.d. exp(1) random variables.

**Remark 2.9** Despite the apparent elegance, it is not straightforward to derive a Central Limit Theorem (CLT) or limit statements concerning W_n_ from the representation of Eq. (3). Initially one could hope (based on “typical” results on limits for randomly weighted sums, e.g. Thm. 1 of Rosalsky and Sreehari, 1998) that W_n_ could converge a.s. to a random variable that has the same limiting distribution as 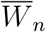.

Similarly, as in the proof of Thm. 2.4 in Section 5, because ((n + 2)(n − 1)/(n(n + 1)) > 1, we have that 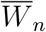 is an L^2^ bounded submartingale

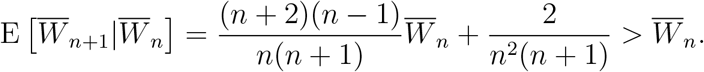

Hence, 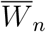 converges almost surely. Figure 1 can easily mislead one to believe in the equality of the limiting distributions of W_n_ and 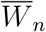. However, in Thm. 6.8 we can see that Var [W_n_] and Var 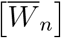 convergence to different limits. Therefore, W_n_ and 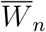 cannot converge in distribution to the same limit. However, as we shall see in Section 4, 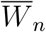 provides a reasonable approximation (and importantly extremely cheap, in terms of computational time and memory) to W_n_ in the sense of their distributions.

**Figure 1:**
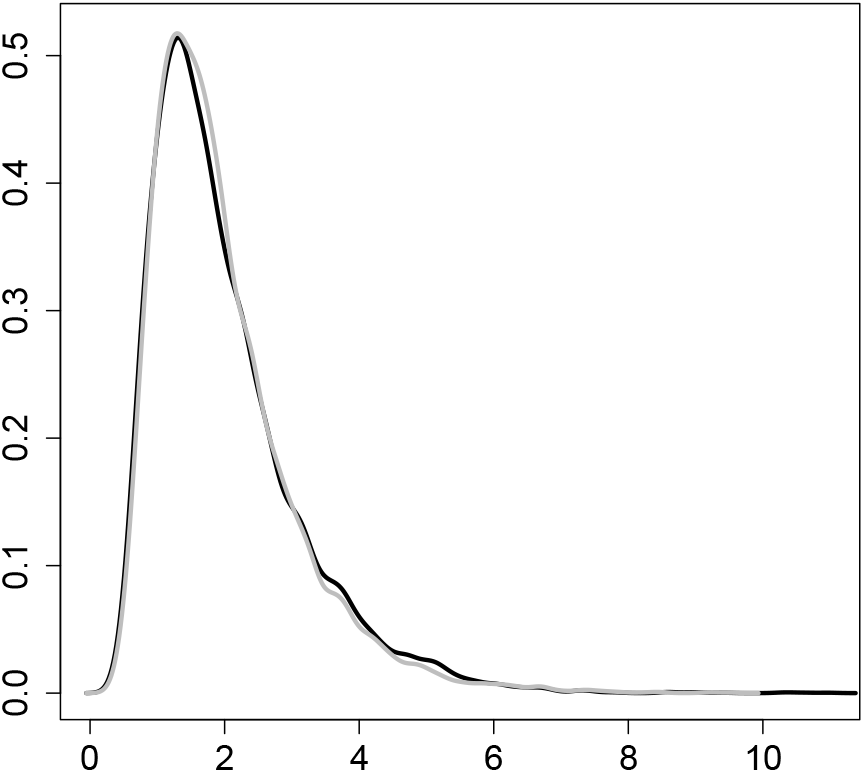
The curves are density estimates, via R’s (R Core Team, 2013) `density()` function, of *W_n_*’s density (black) and 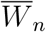’s density (gray). They are based on simulated values of *W_n_* from 10000 simulated 500–tips Yule trees with *λ* =1. To obtain a sample from 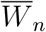, independent exp(1) random variables were drawn. The simulated sample of *W_n_* has mean 2, variance 1.214, skewness 1.609 and excess kurtosis 4.237 while the simulated sample of 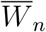 has mean 1.973, variance 1.109, skewness 1.634 and excess kurtosis 4.159. It is obvious that 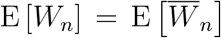, but we have shown that their variances differ (simulations agree with Thm. 6.8).

### Definition 2.10

We naturally define the scaled discrete cophenetic index as

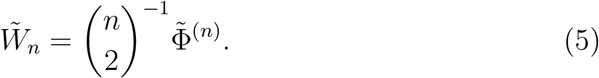

### Theorem 2.11

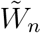 is an almost surely and L^2^ convergent submartingale.

The applied reader will be most interested in how the results here can be practically used. As written already in the Introduction balance indices are often used to provide a single–number summary of the tree’s shape. Such statistics can be then used e.g. to test if the tree is consistent with some null model (here the Yule model). Naturally, there has been extensive work on using different balance indices for significance testing (e.g. Agapow and Purvis, 2002; Blum and François, 2005; Yang et al., 2017). However previous works nearly always worked with indices that only considered the topology and often obtained the rejection regions through direct simulations.

Unfortunately, looking only at the tree’s topology will not allow for distinguishing between some models. In particular (as seen in Tab. 3) there is no difference (from the topological indices perspective) between a Yule tree, a constant rate birth–death tree and a coalescent tree. Hence, a temporal index that also takes into account the branch lengths should be used (as indicated in the “Work remaining” section at the end of Ch. 33 in Felsenstein, 2004). A statistic based on Φ^(*n*)^ performs significantly better (but in these cases still leaves a lot to be desired). However, Φ^(*n*)^ shows it true usefulness when employed to distinguish a biased speciation (Blum and François, 2005) from a Yule model. Blum and François (2005) indicated that there is a regime where topological indices fail completely. Table 3 shows that in this setup (and also certain others) the temporal index in superior in recognizing the deviation from the Yule tree.

Directly simulating a tree from a null model (Yule here) and then calculating the index will of course give a sample from the correct null distribution. However, this approach is costly both in terms of time and memory. Therefore, if theoretical results that provide equivalent, asymptotic or approximate representations of the index’s law are available they could speed up any study by orders of magnitude. In fact this is clearly visible in Tab. 1, calculating the cophenetic index directly from a sample of simulated pure–birth trees is over 170 times slower than considering 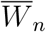. Even more dramatically one can obtain a sample from an approximation to the equivalent representation of the asymptotic distribution of 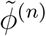 (after normalization) nearly 3000 times faster than directly sampling the discrete cophenetic index.

**Table 1:**
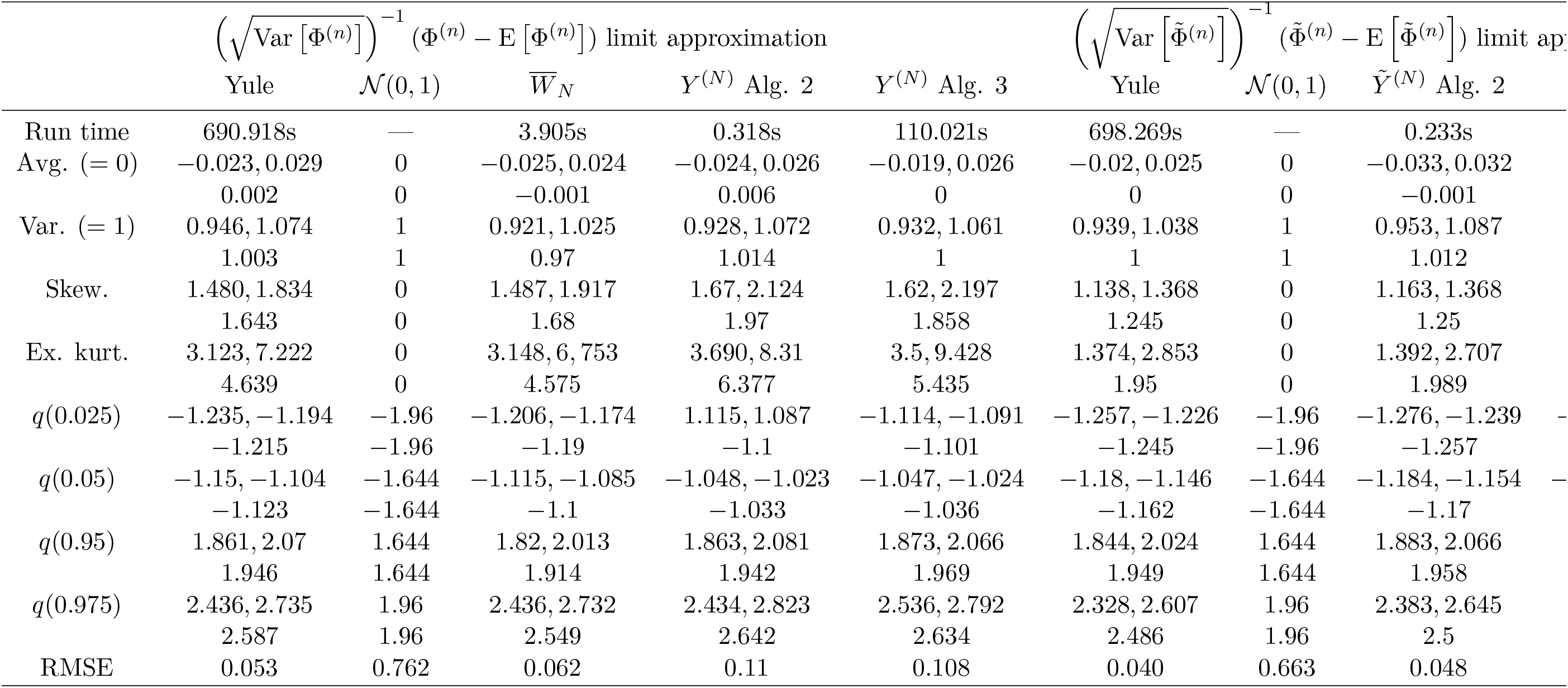
Simulations based on 100 independent repeats of 10000 independent draws of each random variable (population size for Alg. 2) i.e. columns, bar 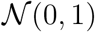. The value on the left is the minimum observed from the 100 repeats, on the right the maximum and in the line below from pooling all repeats together. The running times are averages of 100 independent repeats with 10000 draws each. The abbreviations in the row names are for average (Avg.), variance (Var.), skewness (Skew.) excess kurtosis (Ex. kurt.) and root–mean–square–error (RMSE). The rows *q*(*α*) correspond to the, simulated, bar 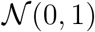, quantiles i.e. for a random variable *X, P*(*X* ≤ *q*(*α*)) = *α*. All simulations were done in R with the package TreeSim (Stadler, 2009, 2011) used to obtain the Yule trees with speciation rate *λ* = 1, *n* = 500 tips and a root edge. The Yule tree φ(^*n*^), 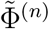 values are centred and scaled by expectation and standard deviation from Eqs. (6) and (7). Other centrings and scalings are summarized in Tab. 2. *N* = 10 for Algs. 2 and 3 is the number of generations and recursion depth of the respective algorithm. In Alg. 2 the initial population is set at 0 and also *Y*_0_ = 0 for Alg. 3. The simulations were run in R 3.4.2 for openSUSE 42.3 (x86_64) on a 3.50GHz. Intel^®^Xeon^®^CPU E5–1620 v4. The calculation of the RMSE is described in the text next to Eq. (15).

In Alg. 1 we describe how the presented here approach can be used for significance testing. Then, in Section 4 we discuss in detail the required computational procedures, present simulation results concerning the power of the tests and apply the tests to empirical data. Preceding this computational Section is a characterization of the limit distribution of (normalized) 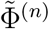 and another proposal to approximate the limit of (normalized) Φ^(*n*)^. This section justifies the described simulation algorithms in Section 4.

**Algorithm 1.**
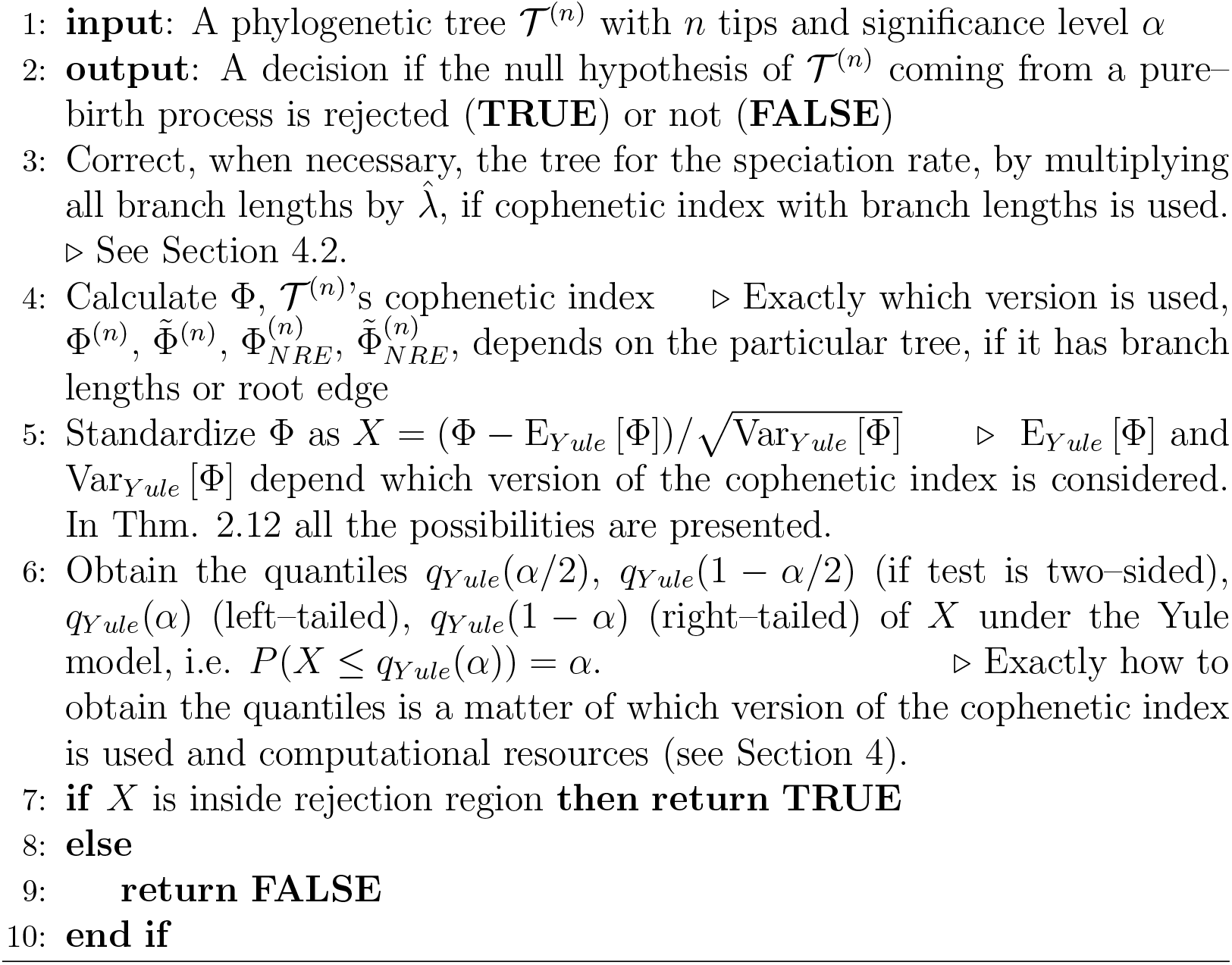
Significance testing

### Theorem 2.12

A random variable with subscript NRE (no root–edge) indicates that this random variable comes from a tree lacking the edge leading to the root.

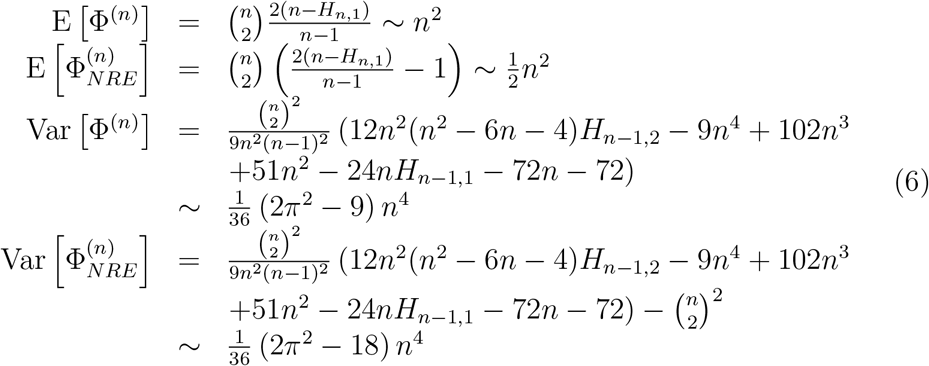

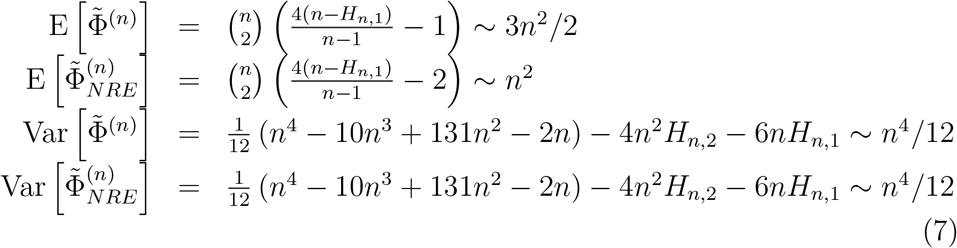

Proof The proof of the expectation part is due to Mir et al. (2013); Sagitov and Bartoszek (2012). The variance of 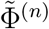 is due to Cardona et al. (2013); Mir et al. (2013). The variance of Φ^(n)^ is a consequence of the lemmata and theorems presented in Section 6. When the root edge is not included, then we have to decrease the expectation by 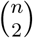. This is due to each pair of tips “having” the root edge included in the cophenetic distance between them. In the case of branch lengths, the expectation of the root edge, distributed as exp(1), is one. Without a root edge for the same reason the variance of Φ^(n)^ has to be decreased by 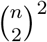. In the discrete case the root edge has a deterministic length of 1 and hence no effect on the variance.

## 3 Contraction–type limit distribution

Even though the representation of Eq. (3) is a very elegant one, it is not obvious how to derive asymptotic properties of the process from it (compare Section 6). We turn to considering the recursive representation proposed by Mir et al. (2013)

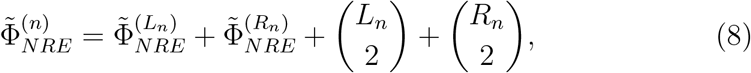

where *L_n_* and *R_n_* are the number of left and right daughter tip descendants. Obviously *L_n_* + *R_n_* = *n*.

From Eq. (8) we will be able to deduce the form of the limit of the process. In the case with branch lengths we attempt to approximate the cophenetic index with the following contraction–type law

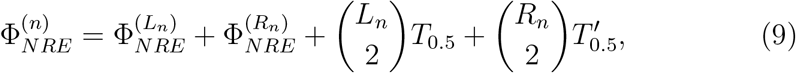

where *T*_0.5_, 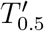 are independent exp(2) random variables (we index with the mean to avoid confusion with *T*_2_, Section 5, the time between the second and third speciation event which is also exp(2) distributed). These are the branch lengths leading from the speciation point. The rationale behind the choice of distribution is that a randomly chosen internal branch of a conditioned Yule tree with rate 1 is exp(2) distributed (Cor. 3.2 and Thm. 3.3 Stadler and Steel, 2012). This is of course an approximation, as we cannot expect that the laws of the branch lengths with the depth of the recursion should become indistinguishable from the law of the average branch. In fact, we should expect that the law of Eq. (9) has to depend on *n*, i.e. the level of the recursion. For larger *n* the branches have distributions concentrated on smaller values, e.g. compare the randomly sampled root adjacent branch length law (Thm. 5.1 Stadler and Steel, 2012) with the law of the average branch length.

However, as we shall see simulations indicate that approximating with the average law still could still yield acceptable heuristics, but not as good as the approximation by 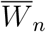. We use the notation 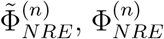 to differentiate from 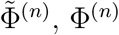 where the root branch is included, i.e.

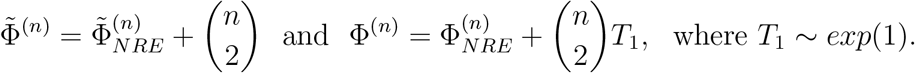

Define now

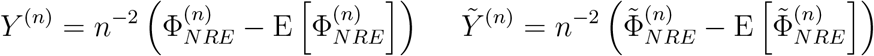

and using Eqs. (6) and (7) we obtain the following recursions

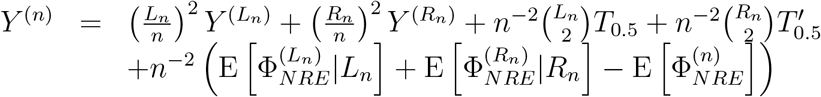

and

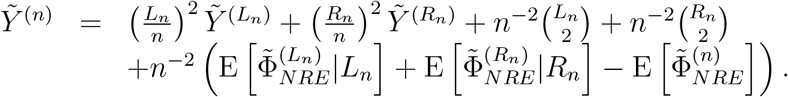

The process *Ỹ*^(*n*)^ is related to the process 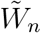 as

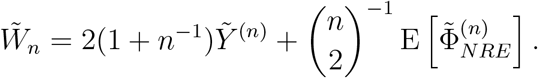

In the continuous case we do not have an exact equality, we rather hope for

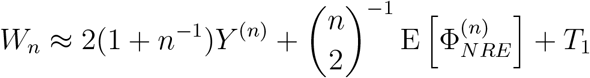

in some sense of approximation. Hence, knowledge of the asymptotic behaviour of *Y*^(∞)^, *Ỹ*^(∞)^ will immediately give us information about *W*^(∞)^, 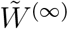 in the obvious way

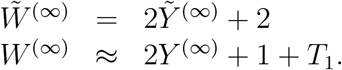

The processes *Y*^(*n*)^, *Ỹ*^(*n*)^ look very similar to the scaled recursive representation of the Quicksort algorithm (e.g. Rösler, 1991). In fact, it is of interest that, just as in the present work, a martingale proof first showed convergence of Quicksort (Régnier, 1989), but then a recursive approach is required to show properties of the limit. The random variable *L_n_*/*n* → *τ* ~ Unif[0, 1] weakly and as weak convergence is preserved under continuous transformations (Thm. 18, p. 316 Grimmett and Stirzaker, 2009) we will have (*L_n_*/*n*)^2^ → *τ*^2^ weakly. Therefore, we would expect the almost sure limits to satisfy the following equalities in distribution (remembering the asymptotic behaviour of the expectations)

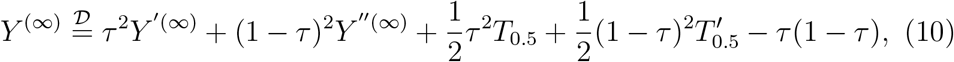

and

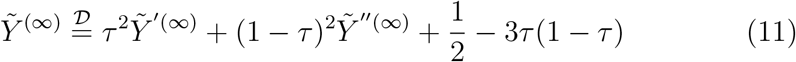

where *τ* is uniformly distributed on [0, 1], *Y*^(∞)^, *Y*^′(∞)^ and *Y*^″(∞)^ are identically distributed random variables, so are *Ỹ*^(∞)^, *Ỹ*^′(∞)^ and *Ỹ*^″(∞)^, and *Ỹ*^′(∞)^, *Y*^″(∞)^, *Ỹ*^′(∞)^ and *∞*^″(∞)^ are independent. Following Rösler (1991)’s approach it turns out that the limiting distributions do satisfy the equalities of Eqs. (10) and (11).

Let *D* be the space of distributions with zero first moment and finite second moment. We consider on *D* the Wasserstein metric

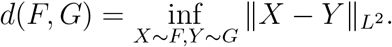

### Theorem 3.1

Let F ∈ D and assume that Y, Y′ ~ F, τ ~ Unif[0, 1], T_0.5_, 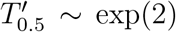 and Y, Y’, τ, T, T’ are all independent. Define transformations S_1_: D → D, S_2_: D → D as

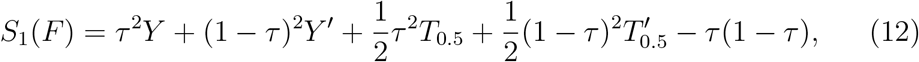

and

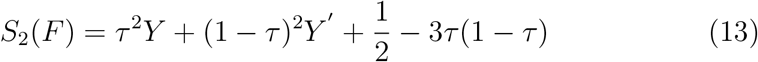

respectively. Both transformations S_1_ and S_2_ are contractions on (D, d) and converge exponentially fast in the d–metric to the unique fixed points of S_1_ and S_2_ respectively.

**Remark 3.2** The proof of Thm. 3.1 is the same as Rösler (1991)’s proof of his Thm. 2.1. However, compared to the Quicksort algorithm (Rösler, 1991) we will have a 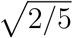 upper bound on the rate of decay instead of 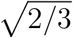. This speed–up should be expected as we have τ^2^ and (1 − τ)^2^ instead of τ and (1 − τ). Thm. 3.1 can also be seen as a consequence of Rösler (1992)’s more general Thms. 3 and 4. The rate of convergence is also a consequence of the general contraction lemma (Lemma 1, Rösler and Rüschendorf, 2001).

Now, using Lemmata 7.1, 7.2 (their proofs in 7.2 differ only in detail from the proof of Prop. 3.2 in Rösler, 1991) and arguing in the same way as Rösler (1991) did in his Section 3, especially his proof of his Thm. 3.1 we obtain that *Y*^(*n*)^ and *Ỹ*^(*n*)^ converge in the Wasserstein *d*–metric to *Y*^(∞)^ and *Ỹ*^(∞)^ whose laws are fixed points of *S*_1_ and *S*_2_ respectively. A minor point should be made. Here, we will have (*i*/*n*)^4^ instead of (*i*/*n*)^2^ in a counterpart of Rösler (1991)’s Prop 3.3.

### Remark 3.3

One may directly obtain from the recursive representation that E [Y^(∞)^] = EỸ^(∞)^ = 0, Var [Y^(∞)^] = 1/16 = 0.0635 and Var Ỹ^(∞)^ = 1/12. We can therefore, see that in the discrete case the variance agrees. However, in the continuous case we can see that it slightly differs

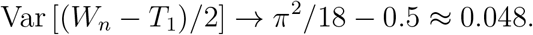

### Remark 3.4

One can of course calculate what the mean and variance of T_0.5_, 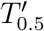 should be so that E [Y^(∞)^] = 0 and Var [Y^(∞)^] = Var [(W_n_ − T_1_)/2]. We should have 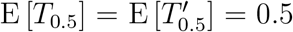 and 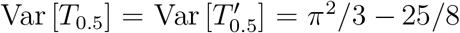. This, in particular, means that these branch lengths cannot be exponentially distributed. We therefore, also experimented by drawing T_0.5_, 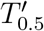 from a gamma distribution with rate equalling 1/(2(π^2^/3 − 25/8)) and shape equalling π^2^/6 − 25/16. However, this significantly increased the duration of the computations but did not result in any visible improvements in comparison to Tab. 1.

## 4 Significance testing

### 4.1 Obtaining the quantiles

Algorithm 1 requires knowledge of the quantiles of the underlying distribution in order to define the rejection region. Unfortunately, an analytical form of the density of any scaled cophenetic index is not known so one will have to resort to some sort of simulations to obtain the critical values. Directly simulating a large number of pure–birth trees can take an overly long time, measured in minutes (on a modern machine with a large amount of memory, or hours on an older one). Fortunately, the cophenetic index can be calculated in *O*(*n*) time (Cor. 3 Mir et al., 2013) and such a tree–traversing algorithm was employed to obtain Φ^(*n*)^ and 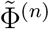. On the other hand, the suggestive (but wrong) approximations of Eq. (4) and contraction limiting distributions Eqs. (10) and (11) are significantly faster to simulate, see Tab. 1.

Simulating from the approximate Eq. (4) is straightforward. One simply draws *n* − 1 independent exp(1) random variables. Simulating random variables satisfying Eqs. (10) and (11) is more involved and it may be possible to develop an exact rejection algorithm (cf. Devroye et al., 2000). Here, we choose simple, approximate but still effective, heuristics in order to demonstrate the usefulness of the approach for significance testing.

We now describe algorithms (Algs. 2 and 3) for simulating from a more general distribution, *F*, that satisfies

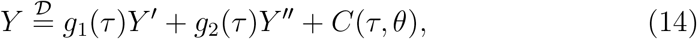

where *Y*, *Y*′, *Y*″ ~ *F, Y*′, *Y*″, *τ,θ* are independent, *τ* ~ *F_τ_, θ* ~ *F_θ_* is some random vector, *g*_1_, *g*_2_: ℝ → ℝ and *C*: ℝ^*p*^ → ℝ for some appropriate *p* that depends on *θ*’s dimension. Of course in our case here we have *τ* ~ Unif[0, 1], *g*_1_(*τ*) = *τ*^2^, *g*_2_(*τ*) = (1 − *τ*)^2^,

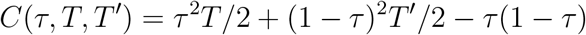

and

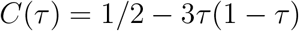

for Φ^(*n*)^, 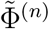 respectively. Of course, *T*, *T*′ are independent and exp(2) distributed. If one considers also the root edge, then to the simulated random variable one needs to add *T*_1_ ~ exp(1) when simulating *n*^−2^Φ^(*n*)^ or appropriately 1 if one considers 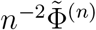.

The recursion of Alg. 3 for a given realization of *τ* and *θ* random variables can be directly solved. However, from numerical experiments implementing Alg. 3 iteratively seemed computationally ineffective.

In Tab. 1 we report on the simulations from the different distribution. For each distribution we draw a sample of size 10000 and repeat this 100 times. We compare the quantiles from the different distributions. We can see that the approximation of 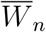 for *W_n_* is a good one and can be used when one needs to work with the distribution of the cophenetic index with branch lengths. In the case of the discrete cophenetic index we have found an exact limit distribution which is a contraction–type distribution. Therefore, one can relatively quickly simulate a sample from it without the need to do lengthy simulations of the whole tree and then calculations of the cophenetic index. Unfortunately, this contraction approach does not seem to give such good results in the Yule tree with branch lengths case. We used an approximation when constructing the contraction. Instead of taking the law of the length of two daughter branches, we took the law of an random internal branch. This induces a difference between the tails of the distributions that is clearly visible in the simulations. Even at the second moment level there is a difference. We calculated (Thm. 6.8) that Var [*W_n_*] → 2*π*^2^/9 − 1 ~ 1.193, 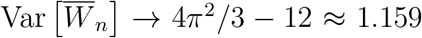 while Var [2*Y*^(*n*)^ + *T*_1_] = 1.25. Therefore, the approximation by 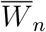 seems better already at the second moment level. Generally if one cannot afford the time and memory to simulate a large sample of Yule tree, simulating 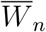 values seems an attractive option, as the discrepancy between the two distributions seems small.

#### Algorithm 2 Population approximation

~~~
1: Initiate population size *N*
2: Set *P*[0, 1 : *N*] = *Y*_0_              [⊲ Initial population
3: **for** *i* = 1 to *i_max_* **do**
4:    *f*_*i*−1_ =density(*P* [*i* − 1,])      ⊲ density estimation by R
5:      **for** *j* = 1 to *N* **d**
6:          Draw *τ* from *F_τ_*
7:          Draw *θ* from *F_θ_*
8:          Draw *Y*_1_, *Y*_2_ independently from *f*_*i*−1_
9:          *P*[*i*,*j*] = *g*_1_(*τ*)*Y*_1_ + *g*_2_(*τ*)*Y*_2_ + *C*(*τ, θ*)
10:   **end for**
11:  **end for**
12:  **return** *P*[*i_max_*,]
13:       ⊲ Add root branch (exp(1) or 1) if needed for each individual.
~~~

#### Algorithm 3 Recursive approximation

~~~
1: **procedure** Yrecursion(*n*, *Y*_0_)
2:   **if** *n* = 0 **then**
3:    *Y*_1_ = *Y*_0_, *Y*_2_ = *Y*_0_
4:   **else if** *n* =1 **and** *Y*_0_ = 0 **then**
5:        Draw *τ*_1_, *τ*_2_ independently from *F_τ_*
6:        Draw *θ*_1_, *θ*_2_ independently from *F_θ_*
7:        *Y*_1_ = *C*(*τ*_1_, *θ*_1_)
8:     *Y*_2_ = *C*(*τ*_2_, *θ*_2_)
9:   **else**
10:     *Y*_1_ =Yrecursion(*n* − 1, *Y*_0_)
11:     *Y*_2_ =Yrecursion(*n* − 1, *Y*_0_)
12:  **end if**
13:    Draw *τ* from *F_τ_*
14:    Draw *θ* from *F_θ_*
15:    **return** *g*_1_(*τ*)**Y**_1_ + *g*_2_ (*τ*)*Y*_2_ + *C*(*τ, θ*),
16: **end procedure**
17: **return** Yrecursion(*N*, *Y*_0_)
18:             ⊲ Add root branch (exp(1) or 1) if needed.
~~~

In Fig. 2 we compare the density estimates of (scaled and centred) both continuous and discrete branches cophenetic indices and their respective contraction–type limit distributions. The density estimates generally agree but we know from Tab. 1 that for Φ^(*n*)^ this is only an approximation. We simulated 10000 Yule trees and hence we report only the quantiles between 2.5% and 97.5%. Quantiles further out in the tails seemed less accurate and hence are not included in the table. Similarly, we can see less correspondence between the different estimates of kurtosis. This statistic relies on fourth moments and hence is more sensitive to the tails. On the other hand we can see much greater Monte Carlo error for the kurtosis in all simulations, including the setup where the values are extracted directly from Yule trees. The values for Φ^(n)^ seem more similar to values from the Yule tree. We should expect this as here we have shown an exact limit distribution.

**Figure 2:**
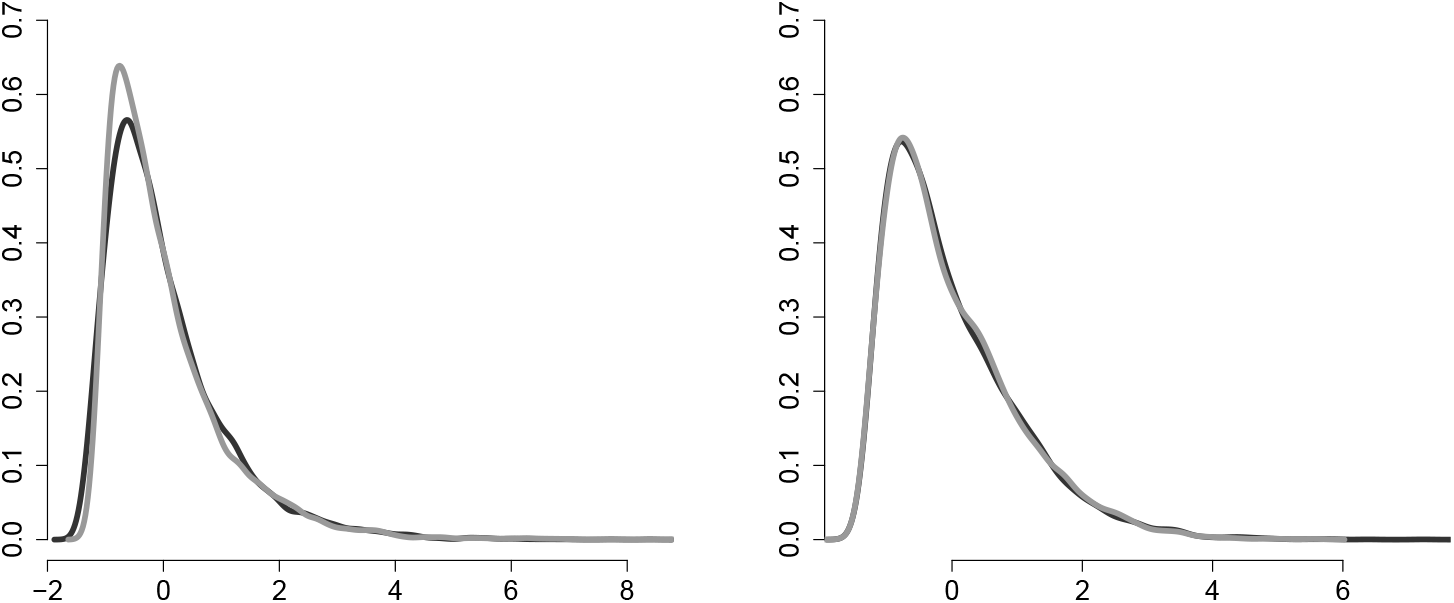
Density estimates of scaled (by theoretical standard deviation) and centred (by theoretical expectation) cophenetic indices (black) from 10000 simulated 500 tip Yule trees with *λ* =1 and of simulation by Alg. 3 (gray), also scaled and centred to mean 0 and variance 1. Left: density estimates for Φ^(*n*)^, right: 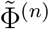. The curves are calculate by R’s `density()` function.

An overall assessment of the quantiles is given by the root–mean–square–error (RMSE) row in Tab. 1. We consider the quantiles at the *α*_1_ = 0.001, *α*_2_ = 0.005, *α*_3_ = 0.01, *α*_4_ = 0.025, *α*_5_ = 0.05, *α*_6_ = 0.95, *α*_7_ = 0.975, *α*_8_ = 0.99, *α*_9_ = 0.995, *α*_10_ = 0.999 levels. The RMSE is defined as

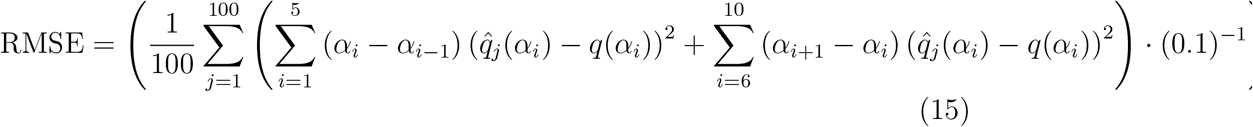

with dummy levels *α*_0_ = 0 and *α*_11_ = 1. The (0.1)^−1^ normalizes the whole mean–square–error. We only look at the error at the tails, so we correct by the fraction of the distributions’ support that we consider. As a proxy for the true quantiles we take the pooled values (as explained in Tab. 1) from the “Yule columns”. The *j* index runs over the 100 repeats of the simulations.

The RMSE, when using 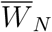, seems to be on the level of the RMSE of the “direct simulations”. *Y*^(*N*)^ has an error of about twice the size (both simulation methods). Looking at *Ỹ*^(*N*)^ one can see that the RMSE is exactly on the level of the “Yule column’s” RMSE. This is even though we used a recursion of level 10, while an exact match of distributions should take place in the limit (infinite depth recursion). However, the rapid, exponential convergence of the contraction seems to make any differences invisible, already at this recursion level.

**Table 2:** Centrings and scalings applied to obtain the entries in Tab. 1. For a random variable X by its centred and scaled version we mean (*X* − *μ*)/*σ*. These centrings and scalings are required to obtain mean zero, variance 1 versions of the random variables, i.e. so that they have the same location and scale as the *z*-transformed cophenetic index. In case of 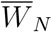 we take the asymptotic scaling (Thm. 6.8) to be comparable with *Y*^(*N*)^. For the convenience of the reader we also provide corresponding centrings and scalings in the no root–edge setup (not considered in Tab. 1).

| | $\overline{W}_N$ | $Y^{(N)}$ | $\tilde{Y}^{(N)}$ |
| --- | --- | --- | --- |
| Centring ($\mu$) | $2(n - H_{n,1})/(n - 1)$ | 1 | 1 |
| Scaling ($\sigma$) | $\sqrt{(2\pi^2 - 9)/9}$ | $\sqrt{1 + 1/16}$ | $(\sqrt{12})^{-1}$ |
| | $\overline{W}_N^{NRE}$ | $Y_{NRE}^{(N)}$ | $\tilde{Y}_{NRE}^{(N)}$ |
| Centring ($\mu$) | $(n + 1 - 2H_{n,1})/(n - 1)$ | 0 | 0 |
| Scaling ($\sigma$) | $\sqrt{(2\pi^2 - 18)/9}$ | 1/4 | $(\sqrt{12})^{-1}$ |

### 4.2 Power of the tests

For a given test statistic to be useful one also needs to know its power, the ability to reject the null hypothesis (here Yule tree) when a given alternative one is true. For example, balance indices based only on topology like Sackin’s, Colless’ or 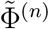 cannot be expected to differentiate between any trees that are generated by different constant rate birth–death processes or by the coalescent. The rationale behind this is that the topologies induced by the n contemporary species (i.e. we forget about lineages leading to extinct ones) are stochastically indistinguishable no matter what the death or birth rate is (Thm. 2.3, Cor. 2.4 Gernhard, 2008). Similarly, regarding the coalescent at the bottom of their p. 93 Steel and McKenzie (2001) write “…, one has the coalescent model [1,18,19]. In this model one starts with n objects, then picks two at random to coalesce, giving *n* − 1 objects. This process is repeated until there is only a single object left. If this process is reversed, starting with one object to give *n* objects, then it is equivalent to the Yule model. Note that in the coalescent model there is commonly a probability distribution for the times of coalescences, but in the Yule model we ignore this element.” To differentiate between such trees one needs to take into consideration the branch lengths. Here we compare the power of the Sackin’s, Colless’, Φ^(*n*)^ and 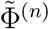 indices at the 5% significance level.

The null hypothesis is always that the tree is generated by a pure–birth process with rate λ =1. The alternative ones are birth–death processes (*λ* = 1, death rate *μ* = 0.25, and 0.5 using the TreeSim package), coalescent process (ape’s `rcoal()` function Paradis et al., 2004) and the biased speciation model for

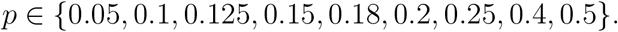

We also simulate a pure–birth process to check if the significance level is met. All trees were simulated with an exp(1) root edge.

The so–called biased speciation model with parameter *p* is the tree growth model as described by Blum and François (2005). In their words, “Assume that the speciation rate of a specific lineage is equal to *r* (0 ≤ *r* ≤ 1). When a species with speciation rate *r* splits, one of its descendant species is given the rate *pr* and the other is given the speciation rate (1 − *p*)*r* where *p* is fixed for the entire tree. These rates are effective until the daughter species themselves speciate. Values of *p* close to 0 or 1 yield very imbalanced trees while values around 0.5 lead to over–balanced phylogenies.” We simulated such trees with in–house R code.

The quantiles of Sackin’s and Colless’ indices were obtained using Alg. 3. It is known (Eqs. 2 and 3 Blum and François, 2005; Blum et al., 2006) that after normalization (centring by expectation and dividing by *n*) in the limit they satisfy a contraction–type distribution of the form of Eq. (14), i.e.

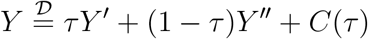

for *τ* ~ Unif[0, 1]. The function *C*(*τ*) takes the form

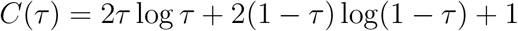

in Sackin’s case and

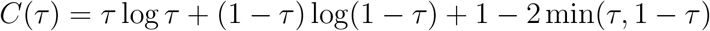

in Colless’ case. It particular, studying the limit of Sackin’s index is equivalent to studying the Quicksort distribution (Blum and François, 2005). We can immediately see the main qualitative difference, the limit of the normalized cophenetic index has the square in *τ*^2^, (1 − *τ*)^2^ in the “recursion part” while Sackin’s and Colless’ have *τ*, (1 − *τ*).

Using 10000 repeats of Alg. 3 with recursion depth 10 we obtained the following sets of quantiles *q*(0.025) = −0.983, *q*(0.95) = 1.189, *q*(0.975) = 1.493, and *q*(0.025) = −1.354, *q*(0.95) = 1.494, *q*(0.975) = 1.868 respectively for the normalized Sackin’s and Colless’ indices.

Under each model we simulated 10000 trees conditioned on 500 contemporary tips. We then checked if the tree was outside the 95% “Yule zone” (Yang et al., 2017) by the procedure described in Alg. 1. We calculated the normalized Sackin’s, Colless’, discrete and continuous cophenetic indices (normalizations from Thm. 2.12). The functions `sackin.phylo()` and `colless.phylo()` of the phyloTop (Kendall et al., 2016) R package were used while the two cophenetic indices were calculated using a linear time in-house R implementation based on traversing the tree (Cor. 3 Mir et al., 2013). Two tests were considered, a two–sided one and a right–tailed one. For the discrete cophenetic index the quantiles from the simulation by Alg. 3 were considered, for the continuous those from 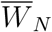 (Tab. 1). The power is then estimated as the fraction of times the null hypothesis was rejected and represented in Tab. 3 by the corresponding Type II error rates. For the Yule tree simulation we can see that the significance level is met. All simulated trees are independent of the trees used to obtain the values in Tab. 1 and quantiles of Sackin’s and Colless’ indices. Hence, they offer a validation of the rejection regions. We summarize the power study in Tab. 3.

**Table 3:**
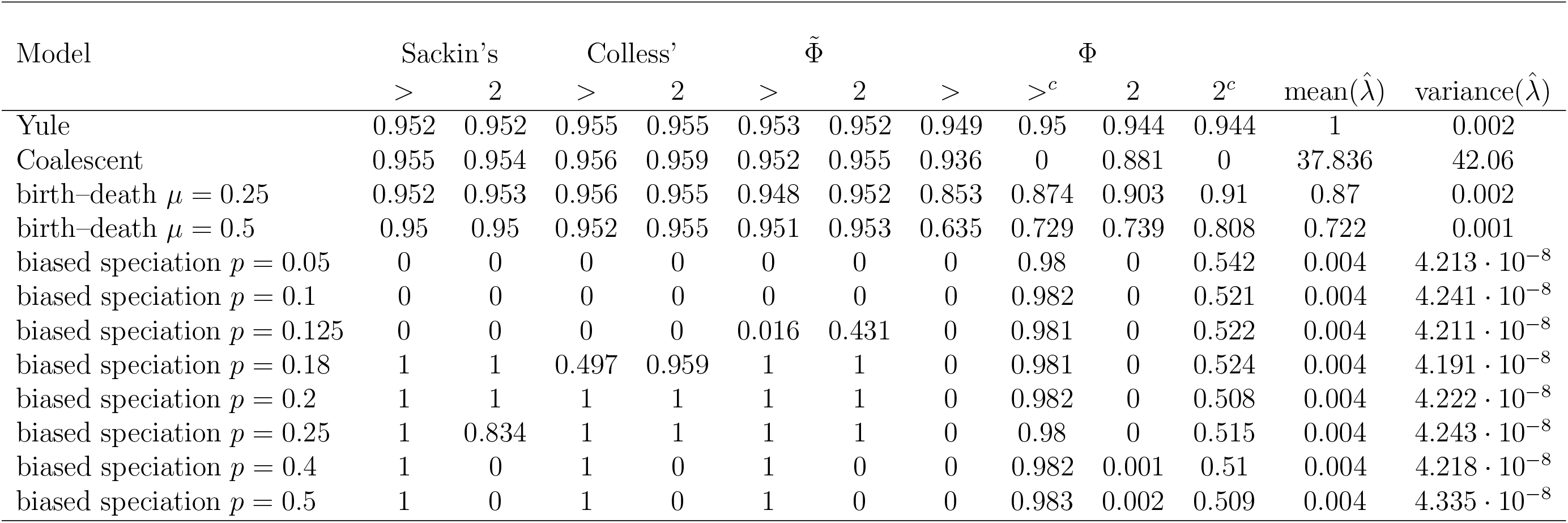
Power, presented as Type II error rates, of the various indices to detect deviations from the Yule tree for various alternative models at the 5% significance level. In the first row the trees were simulates under the Yule (i.e. we present the Type I error rate) so this is a confirmation of correct significance level. Each probability is the fraction of 10000 independently simulated trees that were accepted as Yule trees by the various tests. Columns with “>” label indicate right–tailed test and with label “2” the two–sided test. The critical regions for the cophenetic indices were taken from the pooled estimates in Tab. 1. The superscript *c* indicates tests, where the trees were corrected for the speciation rate through multiplying all branch lengths with 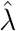. The mean and variance over all trees of 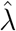, as obtained through ape’s `yule ()` function is reported. Each tree’s branches were scaled by its particular 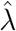 estimate.

As indicated in Alg. 1 one should first “correct” the tree for the speciation rate, when using the cophenetic index with branch lengths. The distributional results derived here on the cophenetic indices are for a unit speciation rate Yule tree. For a mathematical perspective this is not a significant restriction. If one has a pure–birth tree generated by a process with speciation rate *λ* ≠ 1, then multiplying all branch lengths by *λ* will make the tree equivalent to one with unit rate. Hence, all the results presented here are general up to a multiplicative constant. However, from an applied perspective the situation cannot be treated so lightly. For example, if we used the cophenetic index with branch lengths from a Yule tree with a very large speciation rate, then we would expect a significant deviation. However, unless one is interested in deviations from the unit speciation rate Yule tree, this would not be useful. Hence, one needs to correct for this effect. If the tree did come from a Yule process, then an estimate, 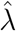, of the speciation rate by maximum likelihood can be obtained. For example, in the work here we used ape’s `yule()` function. Then, one multiplies all branch lengths in the tree by 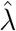 and calculates the cophenetic index for this transformed tree. It is important to point out that 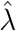 is only an estimate and hence a random variable. The effects of the this source of randomness on the limit distribution deserve a separate, detailed study. Balance indices that do not use branch lengths do not suffer from this issue but on the other hand miss another aspect of the tree—proportions between branch lengths that are non–Yule like.

The power analysis presented in Tab. 3 generally agrees with intuition and the power analysis done by Blum and François (2005). The first row shows that for all tests and statistics the 5% significance level is approximately kept. Then, in the next three rows (coalescent and birth–death process) all topology based indices fail completely (the power is at the significance level). This is completely unsurprising as the after one removes all speciation events (with lineages) leading to extinct species from a birth–death tree, the remaining tree is topologically equivalent to a pure birth tree. The same is true for the coalescent model, its topology is identical in law to the Yule tree’s one. The cophenetic index with branch lengths has a high Type II error rate but is still better, than the topological indices. However, when one “corrects for *λ*” this index manages to nearly (2 trees were not rejected by the two–sided test) perfectly reject the coalescent model trees.

Power for the biased speciation model follows the same pattern as Blum and François (2005) observed. When imbalance is evident, *p* ≤ 0.125, all (*λ* uncorrected) tests were nearly perfect (two–sided discrete cophenetic is an exception). However, the *λ* correction significantly worsened the ability of Φ^(*n*)^ to detect deviations. As imbalance decreased so did the power of the topological indices. For overbalanced trees one–sided tests failed, two sided worked (just as Blum and François, 2005, observed). The cophenetic index with branch lengths (without correction), that does not consider only the topology, was able to successfully reject the pure–birth tree for all p (with only minimal Type II error for *p* ≥ 0.4 in the two–sided test case). Interestingly, Φ^(*n*)^’s (both corrected and uncorrected) power seems invariant with respect to p. These results are especially promising as Φ^(*n*)^ seems to be an index that functions significantly better in the difficult, 0.18 ≤ *p* ≤ 0.25, regime, even after correcting.

At this stage we can point out that a normal approximation to the cophenetic indices’ limit distribution is not appropriate. When doing the above power study we observed that when using the quantiles of the standard normal distribution the right–tailed test based on Φ^(*n*)^ rejects 6.81% of Yule trees, based on 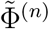 rejects 7.03% of Yule trees, two–sided Φ^(*n*)^ test rejects 4.87% of Yule trees and two–sided 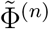 test rejects 4.66% of Yule trees. The Type I error rates of the two–sided tests are within the observed Monte Carlo errors (in Tab. 3) but the right–tailed tests’ Type I error are evidently inflated. This confirms that the right tail of the scaled cophenetic index is much heavier than normal.

In short the power study indicates that the cophenetic index with branch lengths should be considered as an option to detect deviations from the Yule tree. This is because it is able to use information from two sources—the topology and time (a needed direction of development, as indicated in Ch. 33 of Felsenstein, 2004). Actually, this is evident in the decomposition of Eq. (3). The 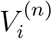s describe the topology and the *Z_i_*s branch lengths. With more information a more powerful testing procedure is possible. Deviations that are not topologically visible, e.g. biased speciation in the 0.18 ≤ *p* ≤ 0.25 regimes, are now detectable. To use Φ^(*n*)^ one should correct for the effects of the speciation rate, as otherwise one merely detects deviations from the unit rate Yule tree. This correction is a mixed blessing. It can help or hinder detection.

### 4.3 Examples with empirical phylogenies

It is naturally interesting to ask how do the indices behave for phylogenies estimated from sequence data. Comparing a database of phylogenies, like TreeBase (http://www.treebase.org), with yet another index’s distribution under the Yule model should not be expected to yield interesting results. The Yule model has been indicated as inadequate to describe the collection of TreeBase’s trees (e.g. Blum and François, 2007). Therefore, we choose a particular study that estimated a tree and also reported a collection of posterior trees. Sosa et al. (2016a) is a recent work, providing all trees from BEAST’s (Drummond and Rambaut, 2007) output, well suited for such a purpose. Sosa et al. (2016a) estimate the evolutionary relationships between a set of 109 tree ferns species. They report a posterior set of 22498 phylogenies (Sosa et al., 2016b).

In Tab. 4 we look what percentage of the trees from the posterior was accepted as being consistent with the Yule tree by the various tests and indices. It can be seen that the discrete cophenetic index has a high acceptance rate. The continuous one, which also takes into account branch lengths did not accept a single tree. However, this is lost when one corrects for the speciation rate (first ape’s `multi2di()` was used to make the trees binary ones). Most tests and indices rejected the Yule tree for the maximum likelihood estimate of the phylogeny with some exceptions. The two–sided discrete cophenetic index test did not reject the null hypothesis of the pure–birth tree. Also after correcting for the speciation rate (estimated at 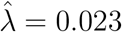), neither test based on the continuous cophenetic index rejected the Yule tree. Therefore, one should conclude (based on the “topological balances”) that the Yule tree null hypothesis can be rejected for this clade of plants.

**Table 4:**
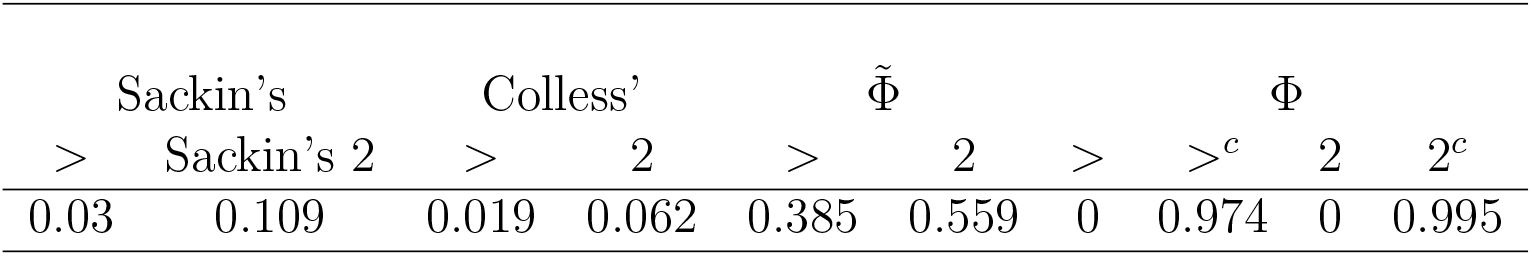
Percentage of trees from Sosa et al. (2016b)’s set of posterior trees accepted as Yule trees by the various tests and indices. Columns with “>” label indicate right–tailed test and with label “2” the two–sided test. The critical regions for the cophenetic indices were taken from the pooled estimates in Tab. 1. The superscript c indicates tests, where the trees were corrected for the speciation rate through multiplying all branch lengths with 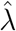. Ape’s `yule()` function returned an average over all trees estimate of *λ* of 0.023 with variance 2.988 · 10^−6^. Each tree’s branches were scaled by its particular λ estimate. Each tree was first transformed by ape’s `multi2di()` into a binary one.

We also followed Blum and François (2005) in looking at Yusim et al. (2001)’s phylogeny of the human immunodeficiency virus type 1 (HIV–1) group M gene sequences, available in the ape R package. The phylogeny consists of 193 tips and Blum and François (2005) could not reject the null hypothesis of the pure–birth tree (using Sackin’s index amongst others). After pruning the tree to keep “only the old internal branches that corresponded to the 30 oldest ancestors” they were able to reject the Yule tree. They conclude that the “results probably indicate a change in the evolutionary rate during the evolution which had more impact on cladogenesis during the early expansion of the virus.” Repeating their experiment we find that only the two versions of the cophenetic index point to a deviation but only in the two–sided test (see Tab. 5). Based only on the 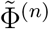’s test and that it conflicts with the conclusions of Sackin’s and Colless’ one should not draw any conclusions. However, as Φ^(*n*)^’s test indicates a deviation, we can be inclined to reject the null hypothesis of the Yule tree. This is further strengthened by the fact that the significance remains after the correction for *λ*. Even though the topology as a whole seems consistent with the pure–birth tree the branch lengths are not. The fact that only the two–sided test rejected the Yule tree indicates that the HIV phylogeny is over–balanced in comparison to a pure–birth tree. In fact, in the biased speciation model tree over–balance is observed for values of *p* close to 0.5 (Blum and François, 2005). Such trees have a declining speciation rate as they grow and hence this supports Blum and François (2005)’s aforementioned explanation.

**Table 5:** Values of the normalized indices for Yusim et al. (2001)’s HIV–1 phylogeny. Above each index is an indication if the index deviates at the 5% significance level from the Yule tree, dash insignificant, asterisk significant. The first symbol concerns the right–tailed test, the second the two–sided test. The superscripted Φ^*c*^ is calculated from the tree corrected for the speciation rate by multiplying all branch lengths by 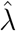.

| Sackin’s | Colless’ | $\tilde{\Phi}$ | $\Phi$ | $\Phi^c$ | $\hat{\lambda}$ |
| --- | --- | --- | --- | --- | --- |
| 0.823<sup>-, -</sup> | 0.993<sup>-, -</sup> | -1.689<sup>-, *</sup> | -1.765<sup>-, *</sup> | -1.602<sup>-, *</sup> | 9.313 |

## 5 Almost sure behaviour of the cophenetic index

We study the asymptotic distributional properties of Φ^(*n*)^ for the pure–birth tree model using techniques from our previous papers on branching Brownian and Ornstein–Uhlenbeck processes (Bartoszek, 2014; Bartoszek and Sagitov, 2015a,b; Sagitov and Bartoszek, 2012). We assume that the speciation rate of the tree is *λ* = 1. The key property we will use is that in the pure–birth tree case the time between two speciation events, *k* and *k* +1 (the first speciation event is at the root), is exp(*k*) distributed, as the minimum of *k* exp(1) random variables. We furthermore, assume that the tree starts with a single species (the origin) that lives for exp(1) time and then splits (the root of the tree) into two species. We consider a conditioned on n contemporary species tree. This conditioning translates into stopping the tree process just before the *n* + 1 speciation event, i.e. the last interspeciation time is exp(*n*) distributed. We introduce the notation that *U*^(*n*)^ is the height of the tree, *τ*^(*n*)^ is the time to coalescent of two randomly selected tip species and *T_k_* is the time between speciation events *k* and *k* + 1 (see Fig. 3 and Bartoszek and Sagitov, 2015b; Sagitov and Bartoszek, 2012).

**Figure 3:**
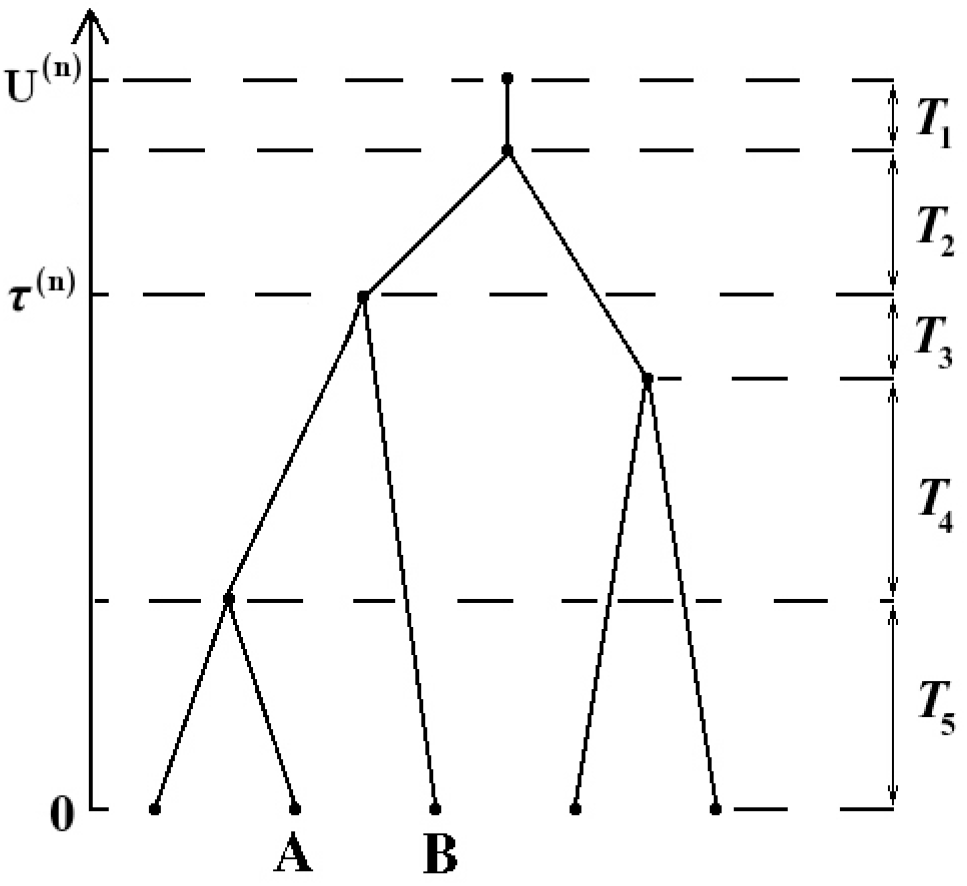
A pure–birth tree with the various time components marked on it. The between speciation times on this lineage are *T*_1_, *T*_2_, *T*_3_ + *T*_4_ and *T*_5_. If we “randomly sample” the pair of extant species “A” and “B”, then the two nodes coalesced at time *τ*^(*n*)^.

### Theorem 5.1

The cophenetic index is an increasing sequence of random variables, Φ^(n+1)^ > Φ^(n)^ and has the recursive representation

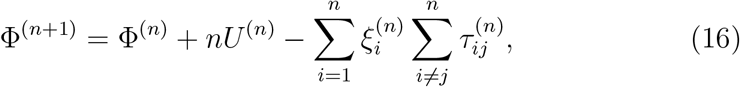

where 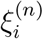 is an indicator random variable whether tip i split at the n–th speciation event.

Proof From the definition we can see that

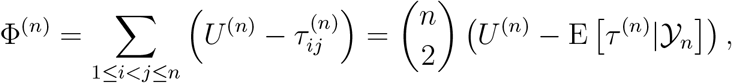

where 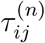 is the time to coalescent of tip species i and j. We now develop a recursive representation for the cophenetic index. First notice that when a new speciation occurs all coalescent times are extended by T_n+1_, i.e.

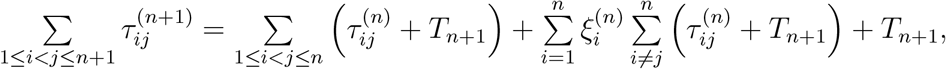

where the “lone” T_n+1_ is the time to coalescent of the two descendants of the split tip. The vector 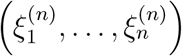 consists of n − 1 0s and exactly one 1 (a categorical distribution with n categories all with equal probability). For each i the marginal probability that 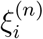 is 1 is 1/n. We rewrite

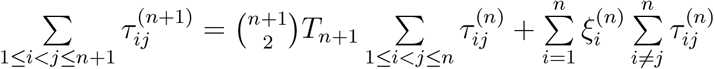

and then obtain the recursive form

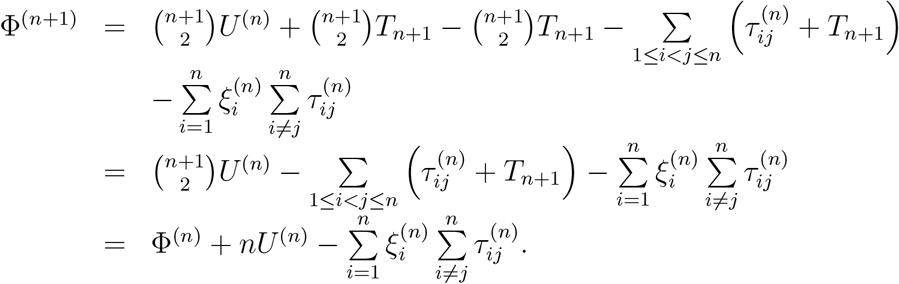

Obviously, Φ^(n+1)^ > Φ^(n)^

Proof of Theorem 2.4. Obviously

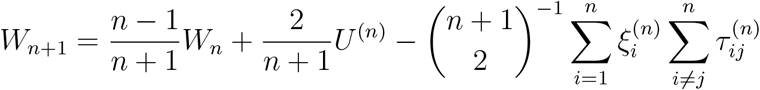

and

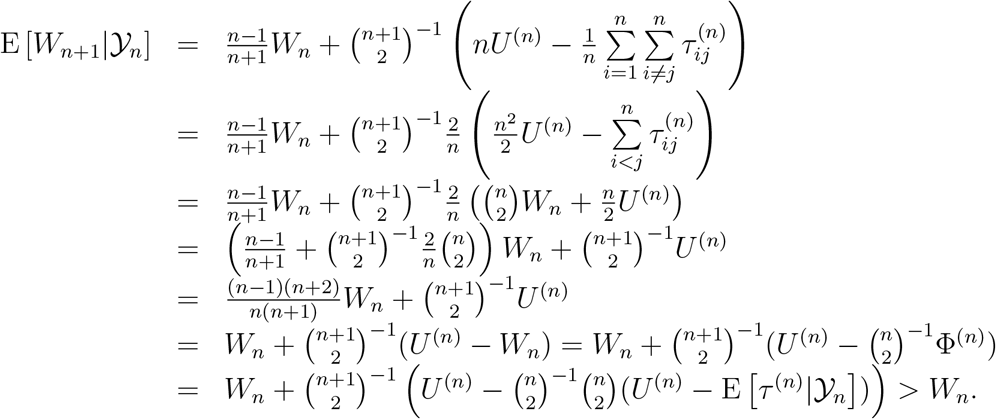

Hence, W_n_ is a positive submartingale with respect to 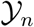. Notice that

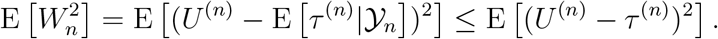

Then, using the general formula for the moments of U^(n)^ − τ^(n)^ (Appendix A, Bartoszek and Sagitov, 2015b), we see that

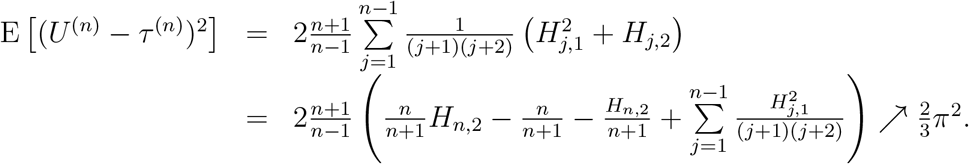

Hence, E[W_n_] and 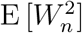 are O(1) and by the martingale convergence theorem W_n_ converges almost surely and in L^2^ to a finite first and second moment random variable.

### Corollary 5.2

W_n_ has finite third moment and is L^3^ convergent.

Proof We first recall the W_n_ is positive. Using the general formula for the moments of U^(n)^ − τ^(n)^ again we see

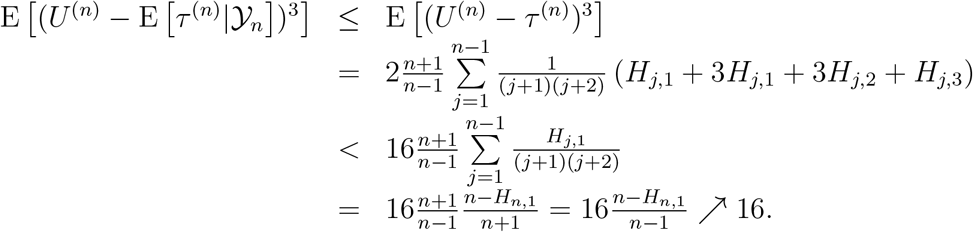

This implies that 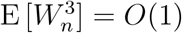 and hence L^3^ convergence and finiteness of the third moment.

### Remark 5.3

Notice that we (Appendix A, Bartoszek and Sagitov, 2015b) made a typo in the general formula for the cross moment of

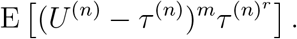

The (−1)^m+r^ should not be there, it will cancel with the (−1)^m+r^ from the derivative of the Laplace transform.

Proof of Theorem 2.7. We write W_n_ as

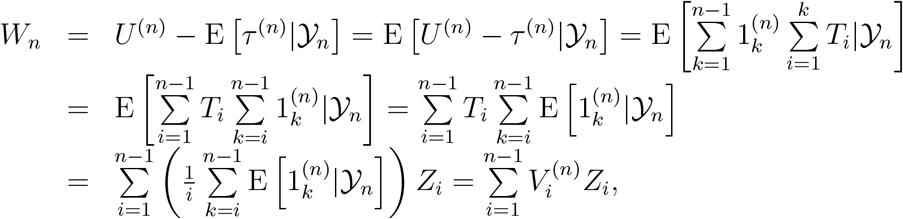

where Z_1_,…, Z_n−1_ are i.i.d. exp(1) random variables.

### Remark 5.4

We notice that we may equivalently rewrite

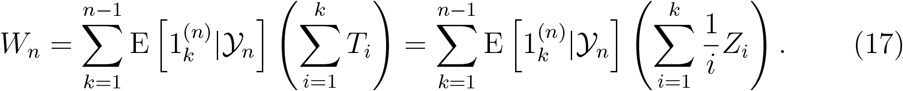

The above and Eq. (3) are very elegant representations of the cophenetic index with branch lengths. They explicitly describe the way the cophenetic index is constructed from a given tree.

Proof of Theorem 2.11. The argumentation is analogous to the proof of Thm. 2.4 by using the recursion

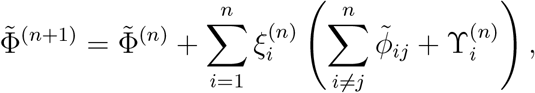

where 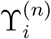 is the number of nodes on the path from the root (or appropriately origin) of the tree to tip i, (see also Bartoszek, 2014, esp. Fig. A.8). An alternative proof for almost sure convergence can be found in Section 7.1.

## 6 Second order properties

In this Section we prove a series of rather technical Lemmata and Theorems concerning the second order properties of 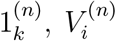 and *W_n_*. Even though we will not obtain any weak limit, the derived properties do give insight on the delicate behaviour of Wn and also show that no “simple” limit, e.g. Eq. (4), is possible. To obtain our results we used Mathematica 9.0 for Linux x86 (64–bit) running on Ubuntu 12.04.5 LTS to evaluate the required sums in closed forms. The Mathematica code is available as an appendix to this paper.

### Lemma 6.1

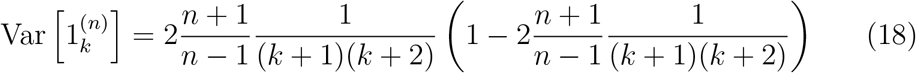

Proof

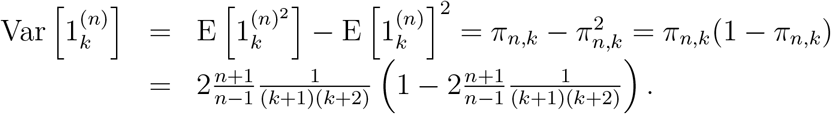

The following lemma is an obvious consequence of the definition of 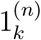.

### Lemma 6.2

For k_1_ ≠ k_2_

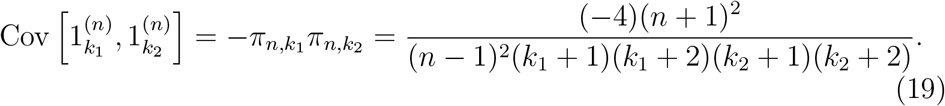

### Lemma 6.3

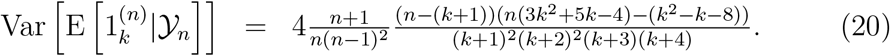

Proof Obviously

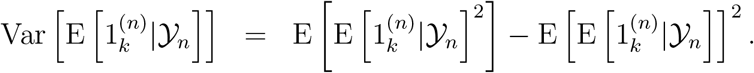

We notice (as Bartoszek and Sagitov, 2015b; Bartoszek, 2016, in Lemmata 11 and 2 respectively) that we may write

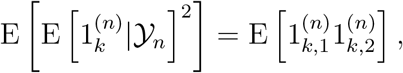

where 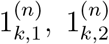 are two independent copies of 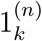, i.e. we sample a pair of tips twice and ask if both pairs coalesced at the k–th speciation event. There are three possibilities, we (i) drew the same pair, (ii) drew two pairs sharing a single node or (iii) drew two disjoint pairs. Event (i) occurs with probability 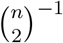, (ii) with probability 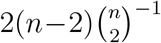 and (iii) with probability 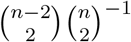. As a check notice that 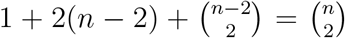. In case (i) 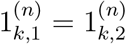, hence writing informally

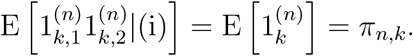

To calculate cases (ii) and (iii) we visualize the situation in Fig. 4 and recall the proof of Bartoszek and Sagitov (2015b)’s Lemma 1. Using Mathe-matica we obtain

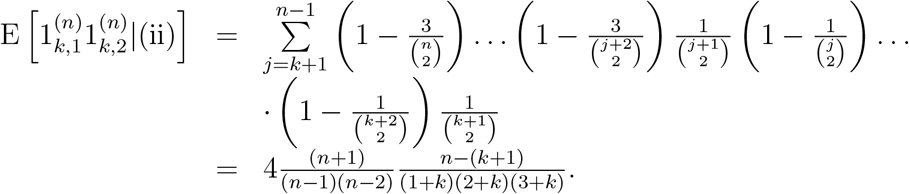

Similarly for case (iii)

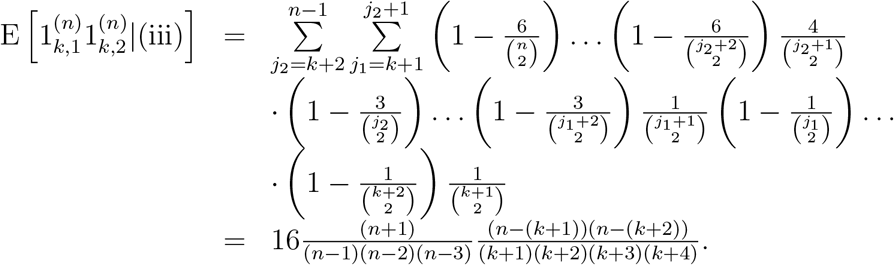

We now put this together as

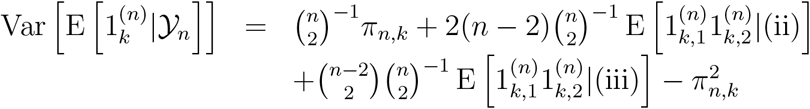

and we obtain (through Mathematica)

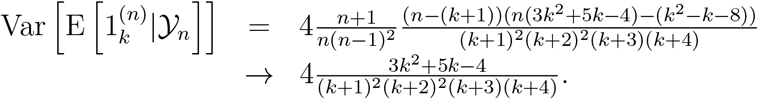

**Figure 4:**
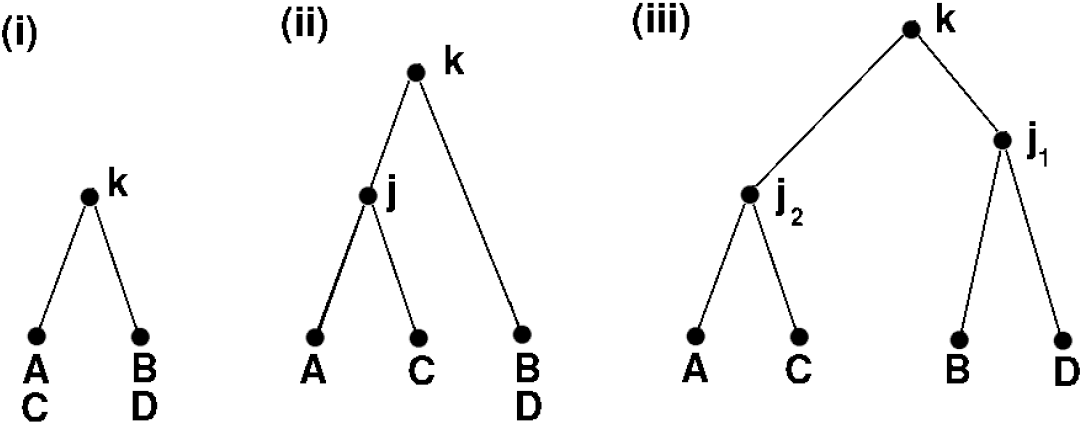
The three possible cases when drawing two random pairs of tip species that coalesce at the *k*–th speciation event. In the picture we “randomly draw” pairs (*A, B*) and (*C, D*).

### Lemma 6.4

For k_1_ < k_2_

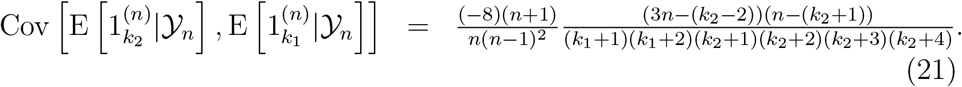

Proof Obviously

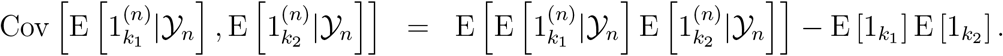

We notice that

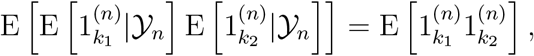

where 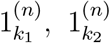 are the indicator variables if two independently sampled pairs coalesced at speciation events k_1_ < k_2_ respectively. There are now two possibilities represented in Fig. 5 (notice that since k_1_ ≠ k_2_ the counterpart of event (i) in Fig. 4 cannot take place). Event (ii) occurs with probability 4/(n +1) and (iii) with probability (n − 3)/(n + 1). Event (iii) can be divided into three “subevents”.

Again we recall the proof of Bartoszek and Sagitov (2015b)’s Lemma 1 and we write informally for (ii) using Mathematica

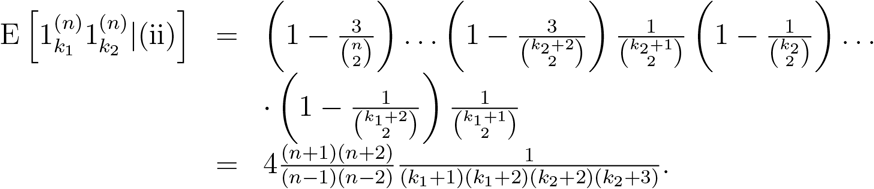

In the same way for the subcases of (iii)

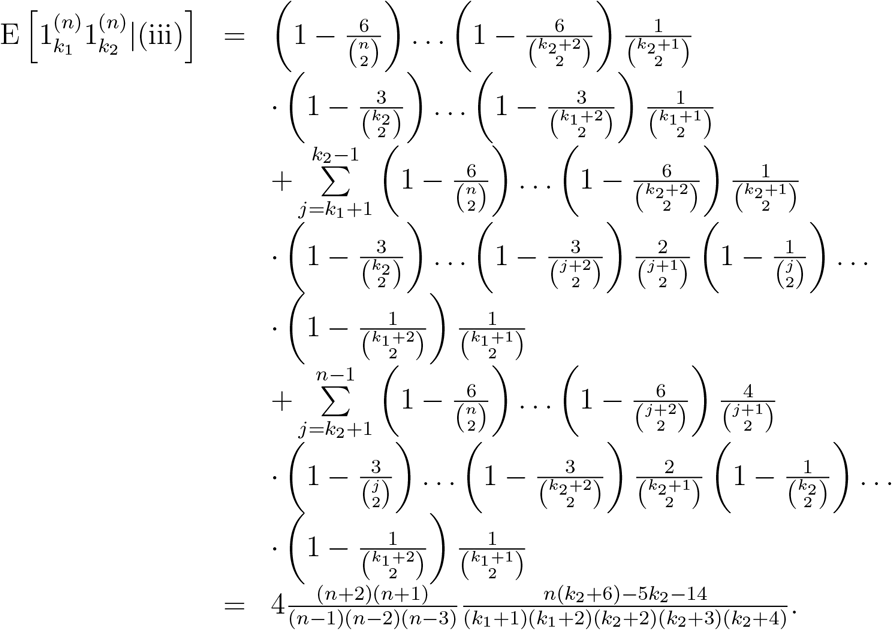

We now put this together as

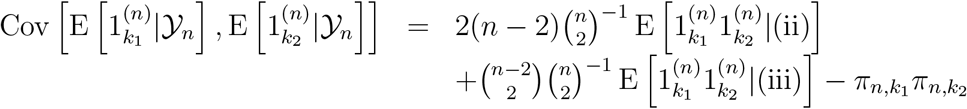

and we obtain

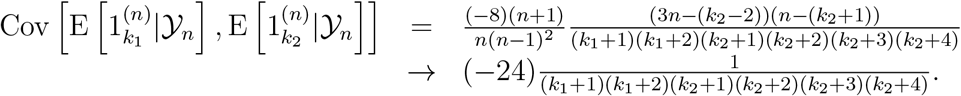

**Figure 5:**
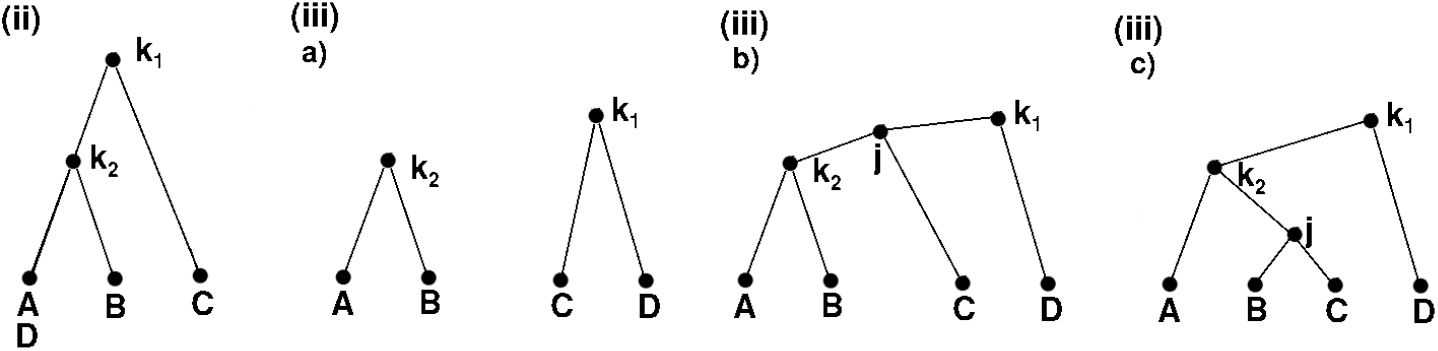
The possible cases when drawing two random pairs of tip species that coalesce at speciation events *k*_1_ < *k*_2_ respectively. In the picture we “randomly draw” pairs (*A, B*) and (*C, D*).

### Theorem 6.5

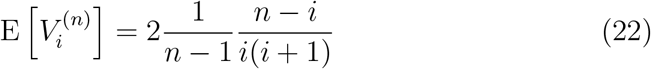

Proof We immediately have

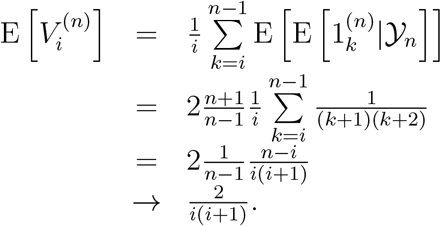

### Theorem 6.6

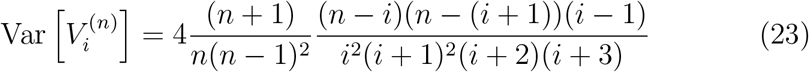

Proof We immediately may write using Lemmata 6.3, 6.4 and Mathematica

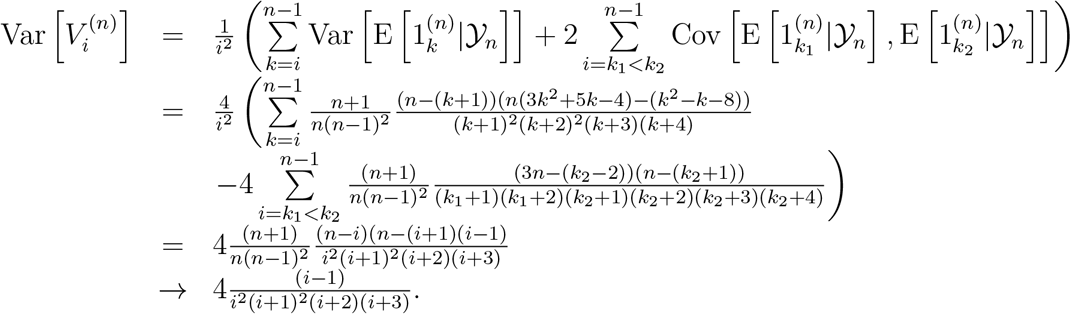

### Theorem 6.7

For 1 ≤ i_1_ ≤ i_2_ ≤ n − 1 we have

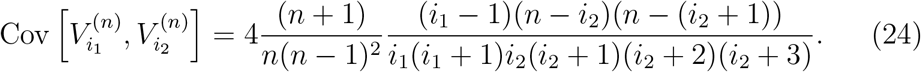

Proof Again using Lemmata 6.3, 6.4, Mathematica and the fact that i_1_ < i_2_

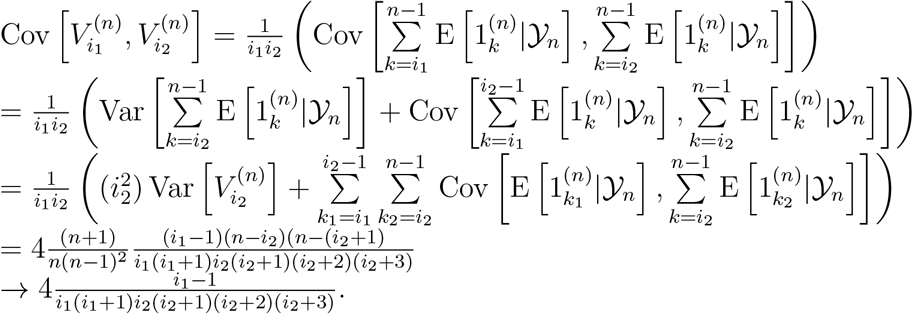

### Theorem 6.8

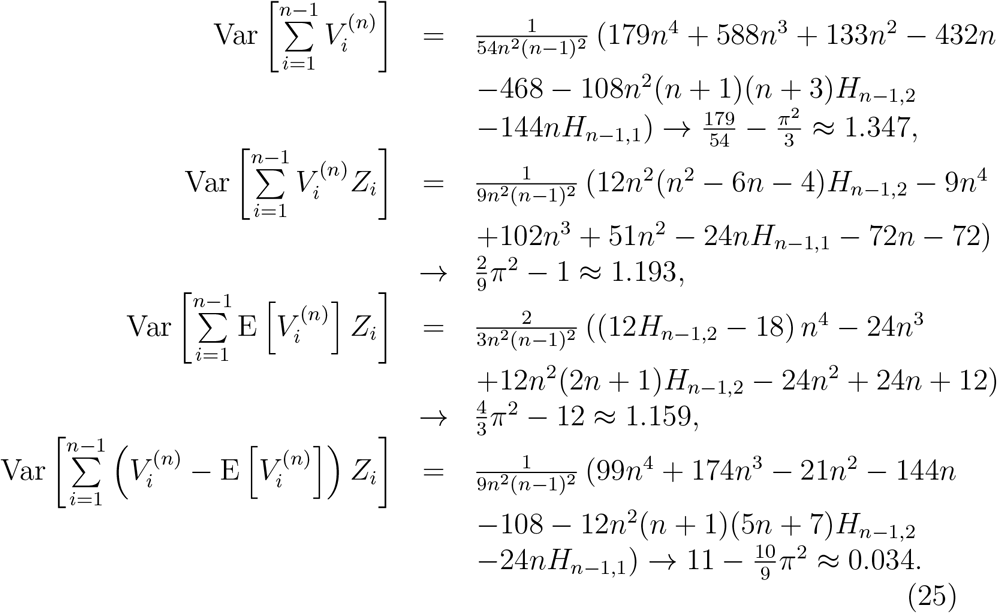

Proof We use Mathematica to first calculate

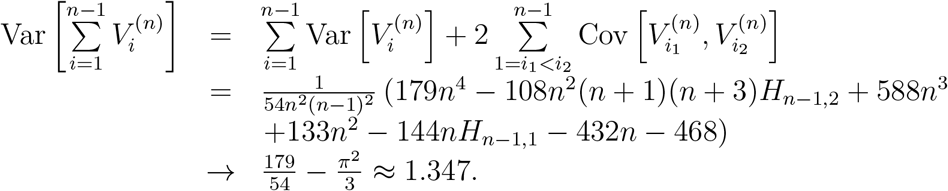

For the second we again use Mathematica and the fact that the Z_i_s are i.i.d. exp(1).

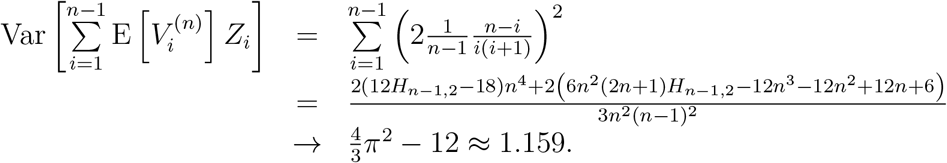

For the third equality we use Mathematica and the fact that for independent families {X} and {Y} of random variables we have

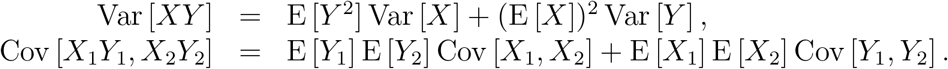

As the Z_i_s are i.i.d. exp(1) we use Mathematica to obtain

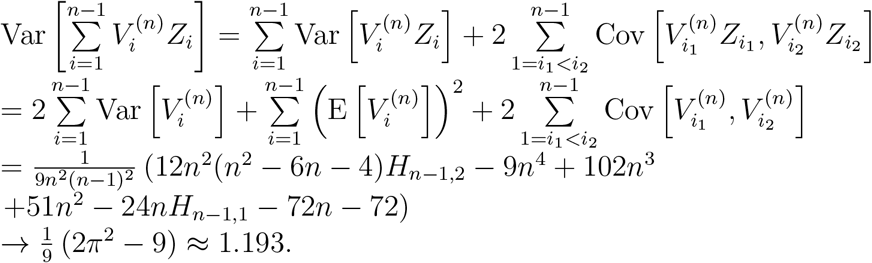

For the fourth equality we use the same properties and pair–wise independence of Z_i_s.

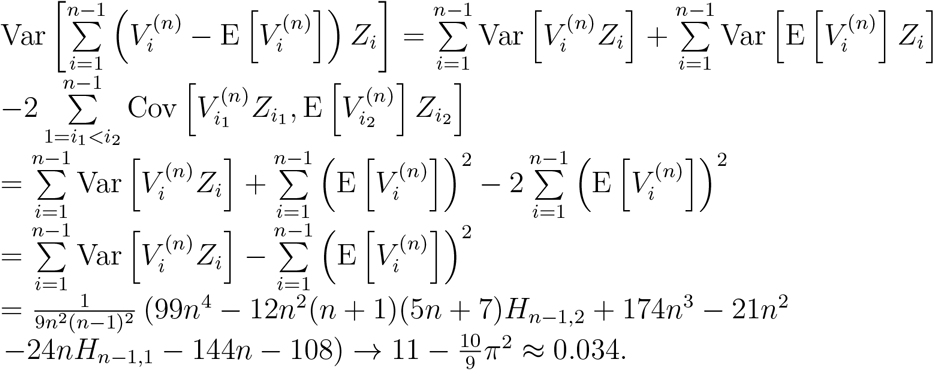

It is worth noting that the above Lemmata and Theorems were confirmed by numerical evaluations of the formulae and comparing these to simulations performed to obtain Fig. 1. As a check also notice that, as implied by variance properties,

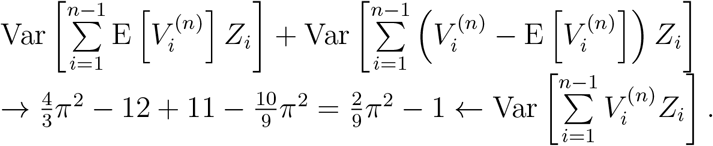

### Theorem 6.9

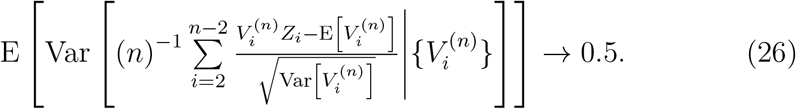

Proof Using the limit for the variance of 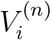 (Thm. 6.6) and the independence of the Z_i_s we have

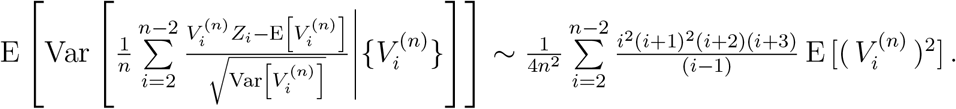

Now from Thms. 6.6 and 6.5 we have

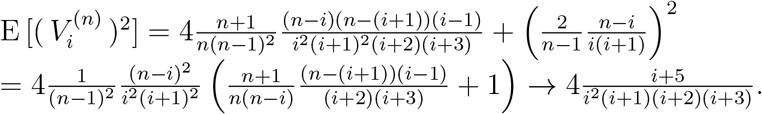

Plugging this in (and using Mathematica)

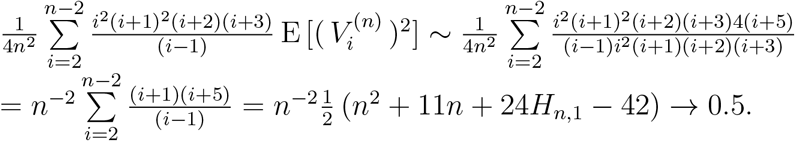

### Remark 6.10

Simulations presented in Fig. 6 and Thm. 6.9 suggest a different possible CLT, namely

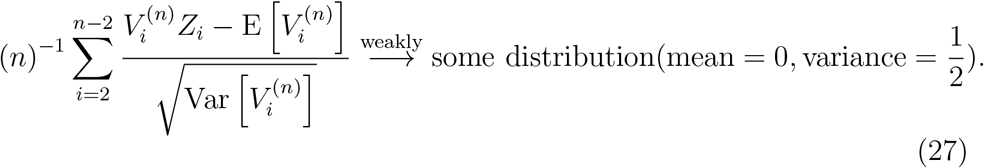

We sum over i = 2,…n − 2 as 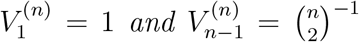 for all n. It would be tempting to take the distribution to be a normal one. However, we should be wary after Rem. 2.9 and Fig. 1 that for our rather delicate problem even very fine simulations can indicate incorrect weak limits. It remains to study the variance of the conditional variance in Eq. (26). It is not entirely clear if this variance of the conditional variance will converge to 0. Hence, it remains an open problem to investigate the conjecture of Eq. (27).

**Figure 6:**
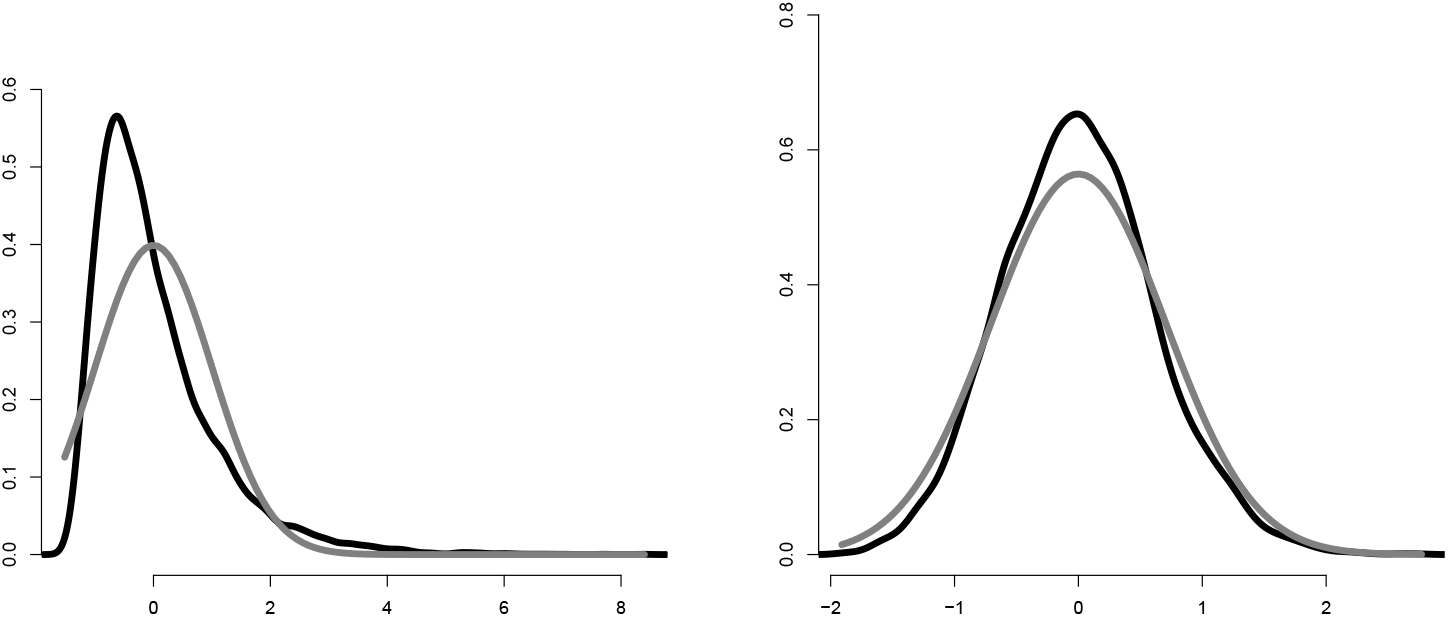
Density estimates of scaled and centred cophenetic indices for 10000 simulated 500 tip Yule trees with *λ* =1. Left: density estimate of 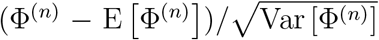. The black curve is the density fitted to simulated data by R’s `density()` function, the gray is the 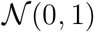 density. Right: simulation of Eq. (27), the gray curve is the 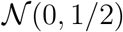 density, and the black curve is the density fitted to simulated data by R’s `density()` function. The sample variance of the simulated Eq. (27) values is 0.385 indicating that with *n* = 500 we still have a high variability or alternatively that the variance of the sample variance in Eq. (26) does not converge to 0.

## 7 Alternative descriptions

### 7.1 Difference process

Let us consider in detail the families of random variables 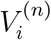 and 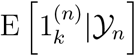. Obviously 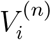 is 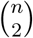 times the number of pairs that coalesced after the *i* − 1 speciation event for a given Yule tree. Denote

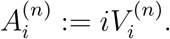

As going from *n* to *n* + 1 means a new speciation event and coalescent at this new *n*th event, then

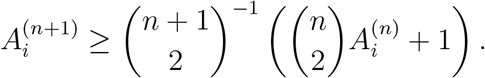

We also know by previous calculations that

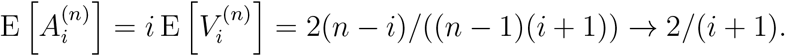

Let 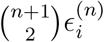 denote the number of newly introduced coalescent events after the (*i* − 1)–one when we go from *n* to *n* +1 species. Obviously 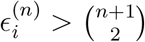. Then, we may write

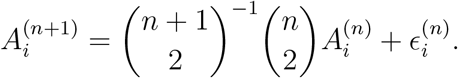

Now,

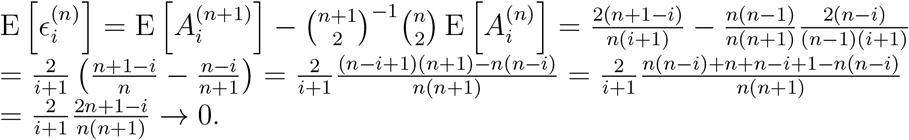

Therefore, for every *i*, 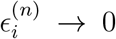 almost surely as it is a positive random variable whose expectation goes to 0. However, 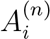 is bounded by 1, as it can be understood in terms of the conditional (on tree) cumulative distribution function for the random variable *κ*—at which speciation event did a random pair of tips coalesce, i.e. for all *i* = 1,…, *n* − 1

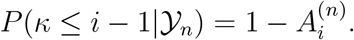

Therefore, as 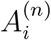 is bounded by 1 and the difference process

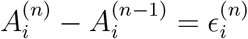

goes almost surely to 0 we may conclude that 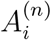 converges almost surely to some random variable *A_i_*. In particular, this implies the almost sure convergence of 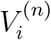 to a limiting random variable *V_i_*. Furthermore, as 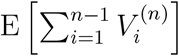 and 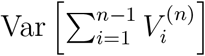 are both *O*(1) we may conclude that 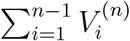 also converges almost surely. This means that the discrete version (all *T_i_* = 1, corresponding to 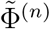) of the cophenetic index converges almost surely (compare with Thm. 2.11).

### 7.2 Polyá urn description

The cophenetic index both in the discrete and continuous version has the following Polyá urn description. We start with an urn filled with *n* balls. Each ball has a number painted on it, 0 initially. At each step we remove a pair of balls, say with numbers *x* and *y* and return a ball with the number (*x* + 1)(*y* + 1) painted on it. We stop when there is only one ball, it will have value 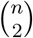. Denote *B_k,i,n_* as the value painted on the *k*–th ball in the *i*–th step when we initially started with *n* balls. Then we can represent the cophenetic index as

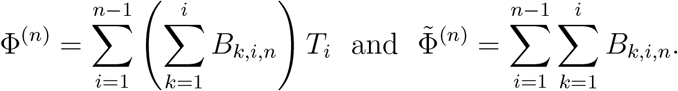

## Acknowledgments

I was supported by the Knut and Alice Wallenberg Foundation and am now by the Swedish Research Council (Vetenskapsrådet) grant no. 2017–04951. I am grateful to the Barcelona Graduate School of Mathematics (BGSMath) for sponsoring the Workshop on Algebraical and Combinatorial Phylogenetics which significantly contributed to the development of my work. I would like to thank the whole Computational Biology and Bioinformatics Research Group of the Balearic Islands University for hosting me on multiple occasions, many discussions and suggestions on phylogenetic indices. My visits to the Balearic Islands University were partially supported by the the G S Magnuson Foundation of the Royal Swedish Academy of Sciences (grants no. MG2015–0055, MG2017–0066) and The Foundation for Scientific Research and Education in Mathematics (SVeFUM). I would like to acknowledge Gabriel Yedid for numerous discussions on the distribution of the cophenetic index and sharing his cophenetic index simulation R code. I am grateful to Cecilia Holmgren and Svante Janson for pointing me to the works on contraction–type distributions and many discussions. I would furthermore like to acknowledge Wojciech Bartoszek, Sergey Bobkov, Joachim Domsta, Serik Sagitov, Mike Steel for helpful comments and discussions related to this work. I am indebted to two anonymous reviewers, an anonymous editor and Haochi Kiang for careful reading of an earlier version of the manuscript and comments significantly improving it.

## Appendix A: Mathematica code for Section 6

(*

Mathematica code used to obtain the closed form formulae of Section 3. Second order properties in K. Bartoszek Exact and approximate limit behaviour of the Yule tree ‘ s cophenetic index.

The script was run using Mathematica 9.0 for Linux x86 (64– bit) running on Ubuntu 12. 04. 5 LTS. It has to be noted that Mathematica ‘ s output should be manually postprocessed in order to have the formulae in terms of harmonic sums and not derivatives of polygamma functions.

All the references in this script point to appropriate fragments of the manuscript.

*)

(*

We choose the pairs in order, i.e. first the first pair to coalesce then the second pair to coalesce.

*)

(* Compare with proof of Lemma 1 of Bartoszek and Sagitov (2015b) *) FcoalProb [ n_, k_, c_]=FullSimplify [Product [(1 − c / ((r * (r − 1)) / 2))

, { r, k + 2,n }]]

(* Def. 2.3, Eq. (1) *)

E1k[n_,k_]: = (2 * (n + 1)/((n − 1) * (k + 1) * (k + 2)))

(* Lemma 6.1, Eq. (18) *)

Var1k[n_,k_]: = (E1k[n,k] −E1k[n,k] * E1k [n,k])

(* Lemma 6.2, Eq. (19) *)

Cov1k11k2 [n_, k1_, k2_]: = (− E1k[n,k1] * E1k [n,k2])

(* Lemma 6.3, Eq. (20) *)

VarE1k [n_,k_] = (FullSimplify [

(1 / (n * (n − 1) /2)) * (2 * (n + 1)/((n − 1) * (k + 1) * (k + 2)))

+

(2 * (n − 2)/(n * (n − 1) / 2)) * (Sum [FcoalProb [n,j,3] * (1 / ((j +1)* j/2))

* FcoalProb[j,k,1] * (1 / ((k+1)* k/2)), {j,k + 1,n − 1}])

+

((n−2) *(n−3)/2/(n *(n−1)/2)) *(

Sum [Sum [ FcoalProb [n,j 1,6] * (4 / ((j 1 +1)* j 1 / 2))

*FcoalProb [j1,j2,3] * (1/((j2+1)*j2/2))

*FcoalProb [j2,k,1] * (1 / ((k + 1) * k/2)), {j1,j2 +1,n − 1}], {j2, k + 1,n − 2 }]

)− (E1k[n,k]) * (E1k[n,k])])

(* Lemma 6.4, Eq. (21) *)

CovE1k1E1k2 [n_,k1_,k2_] = (FullSimplify [

(2 * (n − 2)/(n * (n − 1) / 2)) * (FcoalProb[n,k2,3] * (1/((k2 + 1)* k2 / 2))

*FcoalProb [k2,k1,1] * (1 / ((k1 + 1)* k1 / 2)))

+

((n − 2) * (n − 3)/2/(n * (n − 1) / 2)) * (FcoalProb [n, k2,6]

*(1/((k2 + 1)* k2 /2)) * FcoalProb [k2,k1,3] * (1 / ((k1 + 1)* k1 /2))

+

Sum [FcoalProb [n, k2,6] * (1 / ((k2 + 1)* k2 / 2)) * FcoalProb [k2, j,3]

*(2 / ((j +1)* j / 2)) * FcoalProb [j,k1,1] * (1 / ((k1 + 1)* k1 / 2)), {j, k1 + 1,k2 − 1}]

+

Sum [FcoalProb [n, j,6] * (4 / ((j +1)* j/2)) * FcoalProb [ j, k2, 3]

*(1/((k2 + 1)* k2 / 2)) * FcoalProb [ k2, k1, 1] * (1/((k1 + 1)* k1/2)), {j,k2 + 1,n − 1}]) − (E1k [ n, k1]) * (E1k [ n, k2])])

(* Thm. 6.1, Eq. (2 2) *)

EVi [ n_, i_]: = (FullSimplify [Sum [E1k[n,k], { k,i, n − 1}] / i])

(* Thm. 6.2, Eq. (2 3) *)

Var Vi [ n_, i_] = (FullSimplify [(Sum [ VarE1k [n, k], { k,i, n − 1}]

+2*Sum [Sum [ CovE1k1E1k2 [n,k1,k2], { k2,k1 + 1,n − 1}], { k1, i, n − 1}])/(i * i)])

(* Thm. 6.3, Eq. (2 4) *)

CovVi1 Vi 2 [ n_, i1_, i2_] = (FullSimplify [(i 2 * i 2 * Var Vi [n, i 2]

+Sum [Sum [ CovE1k1E1k2 [n,k1,k2], { k2, i 2, n − 1}], { k1, i 1,i2 − 1}]) / (i 1 * i 2)])

(* Thm. 6.4, formula 1 *)

EVi 2 [n_, i_] = (FullSimplify [ Var Vi [n, i] + (EVi ļn,i]Λ2)])

VarSumVi [n_] = (FullSimplify [Sum [EVi2 [n, i], { i,1, n − 1}]

+2*Sum [Sum [ CovVi1 Vi 2 [n, i 1, i 2], { i2, i 1 +1,n − 1}], { i 1,1, n − 2 }]])

(* Thm. 6.4 Eq. (13), formula 2 *)

VarWn [n_] = (Full Simplify [2 * Sum [ VarVi [n,i], { i,1,n − 1}]

+Sum [ (EVi [n, i])^Λ^2, { i, 1, n − 1}] + 2*Sum [Sum [ CovVi1Vi2 [n, i1, i2]

, { i2, i 1 +1,n − 1}], { i 1,1, n − 1}]])

(* Thm. 6.4 Eq. (13), formula 3 *)

VarWnBar [ n_] = (Full Simplify [Sum [ (EVi [n,i])^Λ^2, { i,1, n − 1}]])

(* Thm. 6.4 Eq. (13), formula 4 *)

VarWnCentre [n_] = (FullSimplify [2 * Sum [VarVi [n,i], { i,1, n − 1}] +2*Sum[Sum[CovVi1Vi2 [n, i 1,i2], { i2, i 1 +1,n − 1}], { i 1,1,n − 1}]])

(* Thm. 6.5 *)

Final Part [ n_] = (Sum [ (i +1) * (i+5)/(i − 1), { i, 2, n − 2 }])

## Appendix B: Counterparts of Rosier (1991)’s Prop. 3.2 for the cophenetic index

### Lemma 7.1

Define for i ∈ {1,…, n}

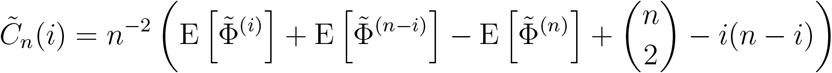

and 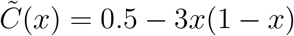 for x ∈ [0, 1], then

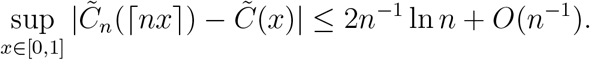

Proof Writing out

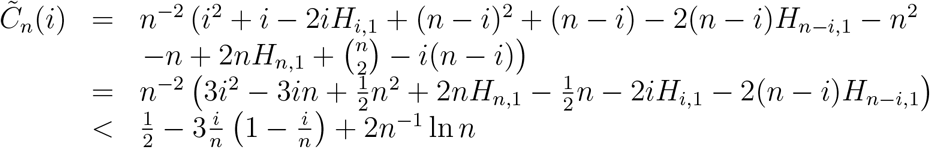

Therefore, assuming that 1 ≤ ⌈nx⌉ ≤ n − 1

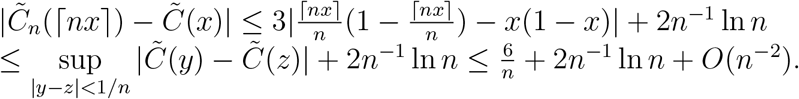

If ⌈nx⌉ = n, we notice that x ∈ (1 − 1/n, 1] and directly obtain

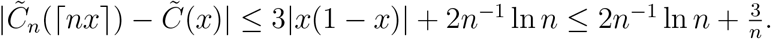

### Lemma 7.2

Define for i ( {1,…, n}, T, T′ ~ exp(2)

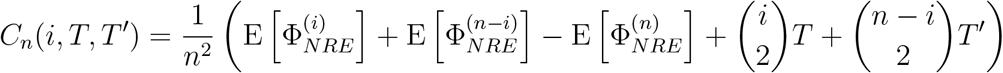

and for x ∈ [0, 1], T, T′ ~ exp(2)

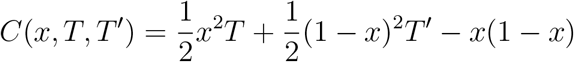

then

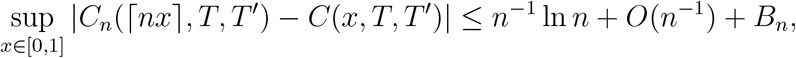

where B_n_ is a positive random variable that converges to 0 almost surely with expectation decaying as O(n^−1^) and second moment as O(n^−2^).

Proof Similarly, as in the proof of Lemma 7.1 we write out

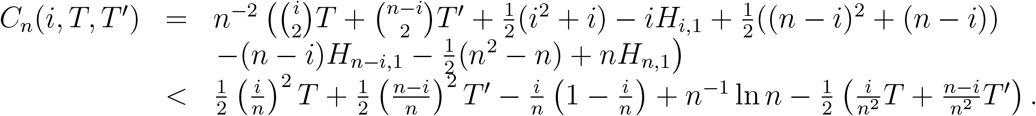

We denote 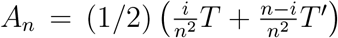 and notice that it converges almost surely to 0 with n. Now, assuming that 1 ≤ ⌈nx⌉ ≤ n − 1

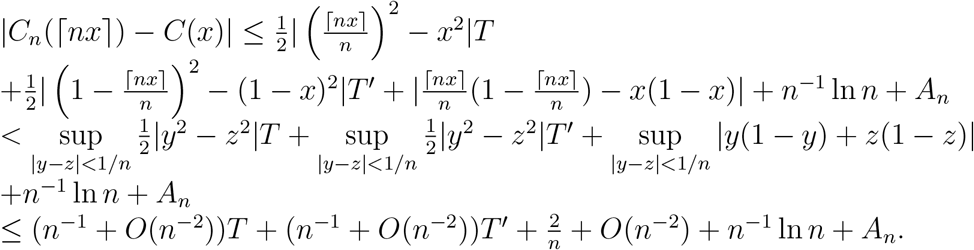

If ∈nx⌉ = n, we notice that x ∈ (1 − 1/n, 1] and directly obtain

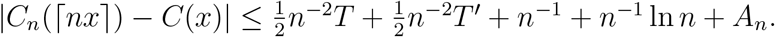

Therefore, if we now denote

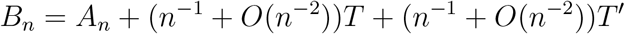

we obtain the statement of the Lemma.

